# DYNAMICS OF RHINOVIRUS SARS-COV-2 COINFECTIONS AND SUPERINFECTIONS IN HUMAN AIRWAY CULTURES REVEAL TYPE-DEPENDENT VIRAL INTERFERENCE

**DOI:** 10.1101/2025.09.24.678304

**Authors:** Romain Volle

## Abstract

Coinfections between respiratory viruses are frequent but their outcomes are poorly understood. Rhinoviruses (RVs) and SARS-CoV-2 are two clinically relevant respiratory pathogens circulating year-round. We used differentiated human bronchial air–liquid interface (ALI) tissue cultures to study coinfections and staggered superinfections between SARS-CoV-2 BA.2 and two RVs (RV-A1, RV-A16). RV-A16 exerted strong and sustained interference on SARS-CoV-2 replication, whereas RV-A1 showed transient effects. SARS-CoV-2 had limited impact on RVs but persisted long-term despite interference. Superinfection demonstrated that pre-established infection with either virus reduced subsequent replication of the other. RNA-FISH revealed spatially distinct infection foci with few dual-infected cells. Although coinfections prolonged interferon and cytokine secretion, functional assays showed that SARS-CoV-2 BA.2 replication was resistant to IFN, in contrast to Wuhan strains. Pleconaril inhibition of RV-A16 spread reduced its interference, highlighting the role of viral spreading. These findings highlight the complexity of respiratory viral interactions and their potential influence on transmission dynamics during viral co-circulation.

## INTRODUCTION

Respiratory tract infections (RTIs) are a major global health burden, ranging from mild upper respiratory illnesses to severe lower respiratory diseases associated with significant morbidity and mortality ^1^. Rhinoviruses (RVs) and coronaviruses (CoVs) are two of the most common causes of RTIs. RVs, with more than 150 known types categorized into three species: RV-A, -B, and -C, are members of the *Picornaviridae* family. RVs are responsible for the majority of common colds but can also trigger severe lower respiratory syndromes ^2,3,4^, and exacerbate chronic airway diseases such as asthma and chronic obstructive pulmonary disease (COPD) ^2,5,6,7,8^. Human coronaviruses typically cause mild upper RTIs, especially the endemic coronaviruses 229E, OC43, HKU1, and NL63. However, the emergence of SARS-CoV in 2003, MERS-CoV in 2013, and most recently SARS-CoV-2 has highlighted their potential for severe acute respiratory disease and pandemic spread. SARS-CoV-2 lead to the COVID-19 pandemic with at least eight reported genetic variants. The last of them, Omicron, continues to expand and diversify manifesting at least 20 phylogenetic clades presently (https://nextstrain.org/ncov/open/global/all-time).

The airway epithelium forms a contiguous barrier and plays an important role for gas exchanges and controlling incoming respiratory pathogens. Airway epithelium is a complex, pseudostratified and heterogeneous tissue with diverse specialized cell populations (*e.g*. Basal, club, goblet, ciliated cells) ^9^. Given that the respiratory tract is a common entry point for pathogens, coinfections are expected to occur relatively frequently. Lung coinfection clinical studies are still scarce, and the underlying mechanisms are poorly understood. A comprehensive quantitative study analyzing >44,000 diagnosed cases of respiratory viral infections over 9-years pointed out highly complex positive and negative interactions between respiratory viruses at the host scale ^10^. Retrospective PCR-based analyses of 1247 nasopharyngeal specimens from symptomatic acute RTIs identified around 10% of specimens with two or more different viruses ^11^. Rhinoviruses were prominently co-detected agents (24%). Rhinoviruses may thus have an unexplored epidemiological regulator function and influence the spread of other respiratory viruses, such as coronaviruses. Experimental studies have reported inconsistent findings on the interactions between RV and SARS-CoV-2 in airways, with results ranging from no effect to significant inhibition of viral replication ^12,13,14^. Furthermore, the mechanisms and long-term outcomes of these interactions remain poorly understood.

Human airway epithelial cultures grown at the air–liquid interface (ALI) closely mimic the architecture and physiology of the bronchial epithelium, providing a relevant model for studying viral coinfections. A key limitation of animal models is that coinfection by two human viruses may be influenced by differences in animal sensitivity, which may not accurately reflect the viral phenotypes in humans. In contrast, tissue culture provides the advantage of testing a wide range of selected experimental parameters at a large scale, enabling the exploration of viral mechanisms with variable factors. Here, we used air-liquid bronchial tissue culture to investigate interactions between SARS-CoV-2 Omicron BA.2 and two RV-A types (RV-A1, RV-A16). We compared simultaneous coinfection and staggered superinfection dynamics, quantified infectious viral titers, assessed innate immune responses, and analyzed spatial distribution of infections. Our findings reveal strong but asymmetric viral interference, with RV-A16 exerting pronounced inhibition of SARS-CoV-2, and point to mechanisms consistent with superinfection exclusion rather than interferon-mediated control.

## RESULTS

### Replication dynamics of coinfecting SARS-CoV-2 (BA.2) and rhinoviruses in bronchial tissue culture

To investigate and compare SARS-CoV-2 (BA.2), RV-A1, and RV-A16 dynamics in both single and coinfection assay, human bronchial air–liquid interface (ALI) cultures from a healthy donor (Epithelix, Geneva; donor #793) were infected with an inoculum of 10,000 TCID₅₀ per virus, corresponding to a theoretical multiplicity of infection (MOI) of ∼0.01 for the entire tissue ^15^. For coinfection, both SARS-CoV-2 (BA.2) and either RV-A1 or –A16 were mixed in the same inoculum. Then, apically released viruses were collected and respectively titrated in parallel on VeroE6-TMPRSS2 cells for SARS-CoV-2 (BA.2) and on HOG cells for RVs (**Fig. 1A**).

**Figure 1.**
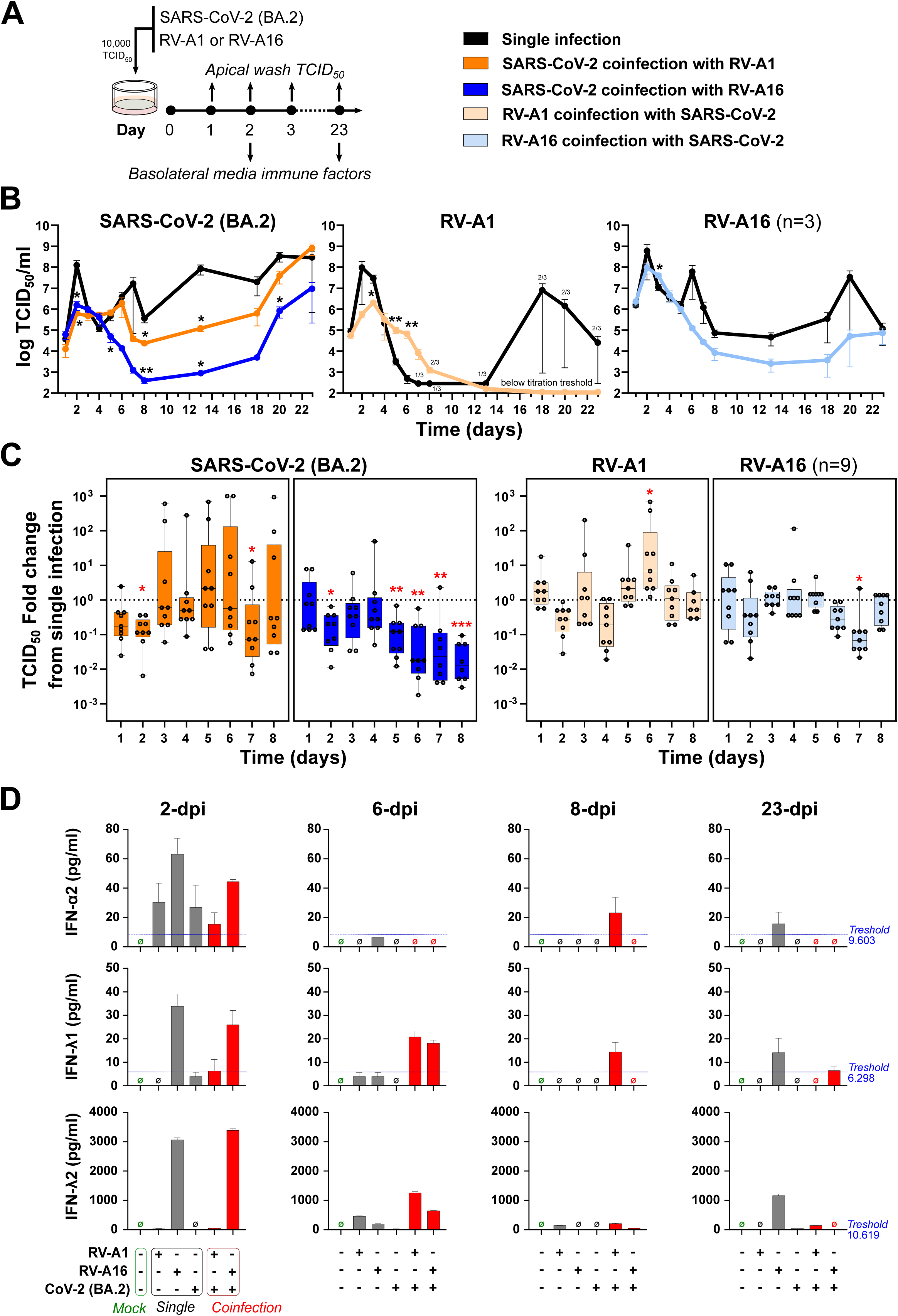
Coinfections of primary bronchial epithelium tissue culture by SARS-CoV-2 (BA.2) and Rhinoviruses. Single donor bronchial epithelium air-liquid cultures were single or simultaneously coinfected by SARS-CoV-2 (BA.2), RV-A1, and RV-A16 at the apical surface (10,000 TCID_50_/tissue, 33.5°C). Virus infectious progenies at the apical surface were collected by apical wash at the indicated time post infections. Apical infectious virus progenies were differentially quantified by TCID_50_ titrations on VeroE6-TMPRSS2 cells for SARS-CoV-2 and Hela Ohio Geneva (HOG) cells for RVs (**A**). Representative TCID_50_ titres kinetic of SARS-CoV-2 (BA.2), RV-A1, and RV-A16 single and coinfections. Data represent the mean of one tissue culture experiment performed in three independent replicate. (**B**). Box-Whisker plots of the virus TCID_50_ titres fold changes between the single infected control (baseline) and the coinfected assay of three independent experiments performed in three independent replicate each (**C**). Interferon concentrations in basolateral media collected at the indicated time post infection were quantified by multiplex microbead-based immunoassay and analyzed. Data are the mean of two independent replicates (**D**). The number of samples with a measurable virus titre is indicated. Statistical comparison was performed using multiple t-test. *P* value for comparison across the groups are shown, *P < 0.05, **P < 0.01, ***P < 0.001.

Following single infection, SARS-CoV-2 (BA.2), RV-A1, and RV-A16 exhibited similar replication kinetics in bronchial tissue cultures, with an acute phase from 1–6 days post-infection (dpi), characterized by peak progeny release at 2–3 dpi, followed by a decline from 3–6 dpi. Notably, SARS-CoV-2 BA.2 and RV-A16 resumed apical release after 5 dpi, whereas most of RV-A1 replicates continued to decline with titres below the detection limit (**Fig. 1B**).

Coinfection experiments revealed a significant reduction of SARS-CoV-2 (BA.2) progeny in the presence of either RV-A1 or RV-A16, most pronounced at the initial replication peak (2 dpi). Importantly, RV-A16 reproducibly (n = 9) maintained significant negative interference on SARS-CoV-2 (BA.2) between 5–8 dpi, whereas RV-A1 exerted only a transient inhibitory effect beyond 2 dpi (**Fig. 1B and 1C**). On the contrary, RV-A1 and RV-A16 progeny release was only mildly reduced by coinfecting SARS-CoV-2 (BA.2) between 1–8 dpi (**Fig. 1B and 1C**).

To assess whether this interference reflected donor-specific effects or a general viral phenotype, bronchial tissue cultures from two additional healthy donors (#839 and #859) were similarly coinfected for 10 days. Despite some donor variability, both tissue cultures showed results consistent with donor #793, with RV-A16 exerting stronger and more sustained interference on SARS-CoV-2 (BA.2) than RV-A1 (**Fig. S1**).

Long-term infections were also examined. As reported in previous work for SARS-CoV-2 (BA.1) ^16^, SARS-CoV-2 (BA.2) and RV-A16 also established persistent infections, with continued production and release of infectious particles for at least 23 days in donor #793 cultures (**Fig. S2**). By contrast, persistent RV-A1 infections were rare. The interference associated by a coinfecting RV-A16 reduced the ability of SARS-CoV-2 to establish long-term infections in 50% of replicates and *vice versa* with RV-A16 persistence. However, persistence of SARS-CoV-2 (BA.2) was also associated with RV-A16 co-persistence, with similar progenies titers as their respective single infection controls (**Fig. S2**). Importantly, tissue integrity, monitored by microscopy and absence of basolateral leakage, remained intact throughout the infection even at viral peaks (2–3 dpi), demonstrating the robust regenerative capacity of the bronchial epithelium model.

Taken together, these findings demonstrate that SARS-CoV-2 (BA.2), RV-A1, and RV-A16 display similar early infection dynamics in human bronchial tissue cultures. However, SARS-CoV-2 replication is strongly modified by coinfecting RVs, with RV-A16 exerting a robust and sustained negative effect, in contrast to the transient inhibition by RV-A1. This effect also extended into long-term infections and was consistent across multiple donors, indicating that RV-A16 acts as a potent interfering agent against SARS-CoV-2 (BA.2) in the bronchial epithelium.

### Innate immune effectors secreted by single and coinfected bronchial tissue culture

Having established that RVs interfere with coinfecting SARS-CoV-2 (BA.2), we next sought to characterize the nature of this interference by analyzing the secretion of innate immune effectors in the basolateral medium. Pools of basolateral samples collected at 2, 6, 8, and 23 days post-infection from single or coinfected tissues (**Fig. 1A**) were analyzed using a multiplex microbead-based immunoassay.

Overall, secretion of soluble immune mediators by the pseudostratified bronchial epithelium varied dynamically according to virus type and infection context. Among interferons, type I IFN-α2 and type III IFN-λ1/λ2 exhibited the highest concentrations (**Fig. 1D**), whereas IFN-β and IFN-γ were present at lower levels but followed a similar secretion pattern (**Fig. S3**). At 2 dpi, IFN responses were the most pronounced in RV-A16 single and coinfected cultures, while SARS-CoV-2 (BA.2) and RV-A1 induced comparatively weak or undetectable IFN secretion. During the virus decline phase (6–8 dpi), IFN levels in single infections approached mock-infected controls, but remained elevated in coinfections (notably SARS-CoV-2 with RV-A1 at 8 dpi). At 23 dpi, low-level secretion of IFN-α2, IFN-λ1, and IFN-λ2 persisted in RV-A16 single infections but not in SARS-CoV-2 (BA.2) and RV-A16 coinfections (**Fig. 1D**).

Several other immune effectors had similar dynamics. These included the immune regulator LIGHT/TNFSF14 ^17^, the pro-survival soluble TNF receptor 2 (TNF-R2) ^18^, the tissue-regenerative IL-10 family (IL-10, IL-20, IL-22, IL-26) ^19,20^, members of the IL-12 family (IL-12p40, IL-27p28, IL-35) ^21,22^, the leukocyte attractants IL-8 and IL-34, the matrix metalloproteinases (MMP-1 to −3), and the inflammation regulator pentraxin-3 ^23^ (**Fig. S3**). At 2 dpi, IL-20 and IL-8 were significantly elevated in RV-A16 single and coinfected cultures but reduced in RV-A1 and SARS-CoV-2 infections, whereas MMP-2 release was increased in all infections, with the highest levels in SARS-CoV-2 coinfections. At 23 dpi, several regulators—including TNFSF14, TNF-R2, IL-10 family cytokines, IL-8, MMP-2, and pentraxin-3—remained elevated in RV-A16 and/or SARS-CoV-2 infections, but not in RV-A1 single infections, consistent with its limited persistence. In contrast, most IL-12 family members and IL-34 returned to baseline, except for IL-27p28, which remained detectable (**Fig. S3**).

Distinct patterns were observed for IL-32, TNFSF12 (TWEAK), TNF-R1, and IL-6–associated pathways. IL-32 ^24^ was consistently elevated in all infections but reached the highest levels in RV-A16 single and coinfected cultures at 2, 8, and 23 dpi. At the opposite, TWEAK/TNFSF12, TNF-R1, and IL-6–associated effectors, including IL-11 ^25^, IL-6Rα, and gp130—were significantly reduced across all infections from 2 dpi and remained suppressed through 23 dpi (**Fig. S3**).

Taken together, these findings indicate that coinfection prolongs for few days the secretion of some innate immune effectors compared with single infections. Moreover, the persistence of RV-A16 and SARS-CoV-2 at 23 dpi was associated with sustained secretion of immune regulators (IL-10, TNFSF14), leukocyte chemoattractants (IL-8), pro-inflammatory mediators (IL-32), and tissue remodeling factors (TNF-R2, MMP-2, IL-22). This prolonged immune activity could be associated with the observed resilience of bronchial tissue cultures, which maintained structural integrity despite ongoing viral replication.

### SARS-CoV-2 (BA.2), RV-A1, and RV-A16 have different sensitivity to IFN

Since RV-A16 was associated with both the strongest interference against SARS-CoV-2 (BA.2) and the highest levels of soluble immune effectors in the basolateral medium, we investigated whether these two phenotypes were functionally linked. Bronchial tissue cultures were infected with RV-A16 or mock-infected for 2 days, as described above, after which basolateral media were collected. RV-A16 infectious particles from the collected conditioned media were inactivated by a treatment with the rhinovirus entry inhibitor Pleconaril (1 µM) ^26^ for 2 h. Apical RV-A16 titration controls confirmed productive infection, yielding a titre of 6.9 ± 0.5 log₁₀ TCID₅₀. Then RV-A16 and mock conditioned basolateral media were used to prime naïve bronchial tissue cultures for 24 h, prior to the infection with 10,000 TCID_50_ of SARS-CoV-2 (BA.2) or SARS-CoV-2 (Wuhan), as control. Apical SARS-CoV-2 titres were quantified by TCID₅₀ assay on both VeroE6-TMPRSS2 cells and HOG cells, as a control of effective RV-A16 inactivation by Pleconaril treatment (**Fig. 2A and 2B**). Apical SARS-CoV-2 titration from tissue cultures primed with RV-A16–conditioned media showed a significant 1-log reduction in TCID₅₀ titres for the Wuhan variant up to 2 dpi compared with mock-primed cultures, whereas no significant differences were observed for SARS-CoV-2 BA.2 (**Fig. 2C and 2D**).

**Figure 2.**
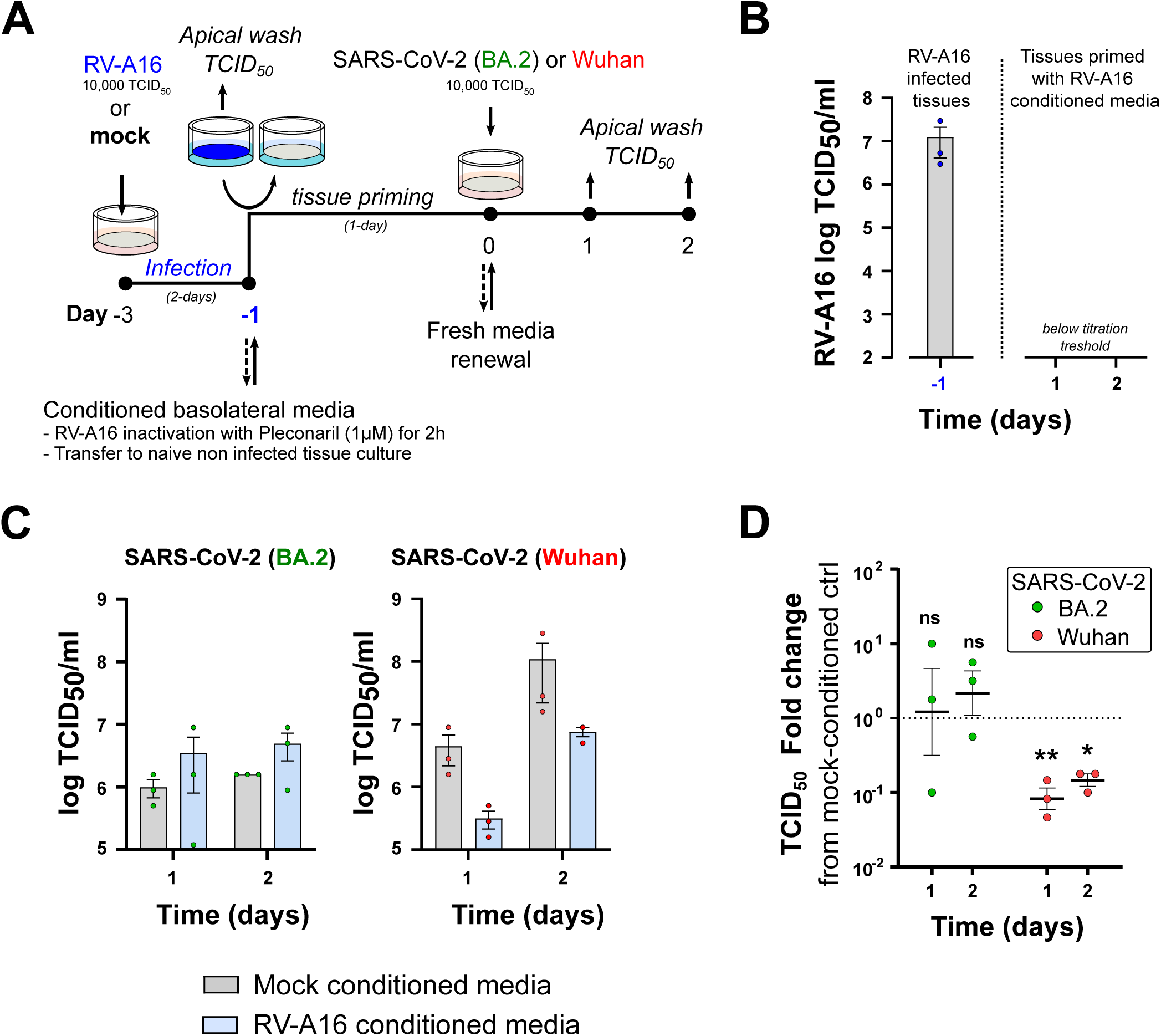
SARS-CoV-2 (BA.2) resistance to RV-A16 conditioned basolateral media. Single donor bronchial epithelium air-liquid cultures were single infected at the apical surface by RV-A16 (10,000 TCID_50_/tissue, 33.5°C) or mock infected for two days. RV-A16 and mock-infected conditioned basolateral media were respectively collected, pooled, and treated for 2-hours by addition of Pleconaril (1µM) for RV-A16 inactivation. Then conditioned basolateral media were used to prime naive non-infected bronchial epithelium air-liquid cultures for 24-hours, prior to be infected by either SARS-CoV-2 BA.2 or Wuhan variants for 2-days. Virus infectious progenies at the apical surface were collected by apical wash at the indicated time post infections (**A**). RV-A16 TCID_50_ titres (**B**) and SARS-CoV-2 TCID_50_ titres (**C**) are represented as one tissue culture experiment performed in three independent replicates. SARS-CoV-2 TCID_50_ titres fold changes between tissue cultures primed with RV-A16 and mock-infected (baseline) conditioned basolateral media (**C**). Data are represented as mean±SEM. Statistical comparison was performed using multiple t-test. *P* value for comparison across the groups are shown, *P < 0.05, **P < 0.01, *ns* for non-significant.

We next tested whether the elevated IFN-α2, IFN-λ1, and IFN-λ2 detected in RV-A16 infections (**Fig. 1D**) contributed to this interference. Bronchial epithelial cultures were primed for 24 h with a mixture of IFN-α (60 pg/mL), IFN-λ2 (3,000 pg/mL), and IFN-λ1 (30 pg/mL) supplemented to their basolateral media. Used concentrations correspond to those measured at 2 dpi in RV-A16-infected tissues. After refreshment of media, cultures were single or coinfected with RV-A1, RV-A16, SARS-CoV-2 (BA.2), or SARS-CoV-2 (Wuhan) (**Fig. 3A**). In agreement with our previous results (**Fig. 2**), Wuhan strain exhibited a significant reduction in titers following IFN priming, whereas SARS-CoV-2 (BA.2) replication was unaffected by IFN treatment in both single and coinfections (**Fig. 3B**). Notably, SARS-CoV-2 (BA.2) coinfected with RV-A1 displayed a transient 1-log reduction at 2 dpi under IFN priming, but titers recovered thereafter. RV-A1 was highly sensitive to IFN priming, with a 2-log reduction in titres at 2 dpi and recovering in titres at 4 dpi in both single and coinfections (**Fig. 3C**), whereas RV-A16 replication was unaffected (**Fig. 3D**).

**Figure 3.**
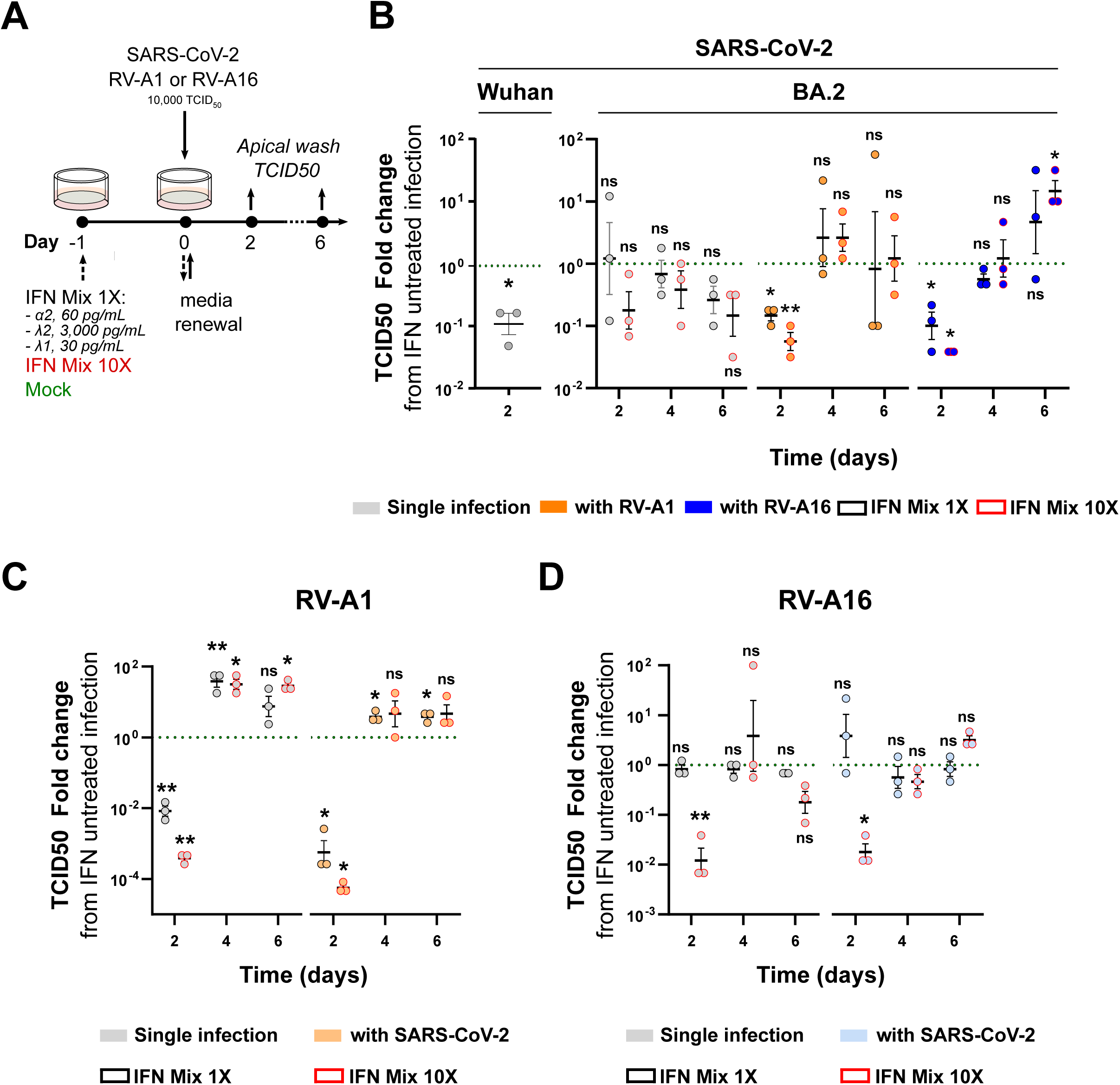
Differential virus sensitivities to IFN pretreatment. Single donor bronchial epithelium tissue cultures were primed for 1-day by addition in the basolateral media of a mix of IFN-α (60 pg/ml), IFN-λ2 (3,000 pg/mL) and IFN-λ1 (30 pg/mL), called IFN Mix 1X (results squared in black); or of a mix of IFN-α (600 pg/ml), IFN-λ2 (30,000 pg/mL) and IFN-λ1 (300 pg/mL), called IFN Mix 10X (results squared in red). Then prior virus inoculation, IFN treatment was removed by basolateral media renewal and epithelia were apically single or simultaneously coinfected by SARS-CoV-2 (BA.2), RV-A1, and/or RV-A16 (10,000 TCID50/tissue, 33.5°C). IFN non-primed tissue culture controls (Mock) were similarly infected in parallels. Virus infectious progenies at the apical side have been differentially quantified at the indicated time points by TCID_50_ titrations. (**A**). Virus titres fold changes between the IFN treated sample and its mock treated corresponding control (baseline) are represented as one tissue culture experiment performed in three independent replicates for SARS-CoV-2 (BA.2) (**B**), RV-A1 (**C**), and RV-A16 (**D**). Bars represents mean±SEM. Statistical comparison was performed using multiple t-test. *P* value for comparison across the groups are shown, *P < 0.05, **P < 0.01, *ns* for non-significant.

To further assess IFN resistance, we repeated the experiment using a 10-times higher IFN mixture with IFN-α (600 pg/mL), IFN-λ2 (30,000 pg/mL), and IFN-λ1 (300 pg/mL). SARS-CoV-2 (BA.2) replication remained largely unaffected in single infection, though coinfections with RV-A1 or RV-A16 showed a 1-log reduction at 2 dpi (**Fig. 3B**). In contrast, both rhinoviruses exhibited strong reduction. RV-A1 titers were reduced by 4 logs and RV-A16 by 2 logs in both single and coinfections (**Fig. 3C and 3D**).

Altogether, these results indicate that type I and III interferons do not play a significant role in the interference of SARS-CoV-2 (BA.2) by coinfecting rhinoviruses. Furthermore, our data highlight a greater sensitivity to IFN of RVs compared to SARS-CoV-2 (BA.2). This particular IFN susceptibility of RV-A1 may contribute to its inability to establish sustained persistent infections in tissue cultures, especially during coinfection with SARS-CoV-2 (BA.2), where elevated IFN levels were observed at 6–8 dpi (**Fig. S3**).

### Distinct and overlapping infection foci of coinfecting SARS-CoV-2 and Rhinoviruses

To assess the spatial distribution and persistence of RVs and SARS-CoV-2, viral RNA fluorescence *in situ* hybridization (FISH) was performed on single- and coinfected bronchial tissue cultures fixed at 8 dpi. Widefield fluorescence microscopy revealed strong and widespread SARS-CoV-2 signals in single infections, whereas RV-A1 and RV-A16 displayed more localized staining. In coinfected cultures, however, both viruses exhibited restricted distributions confined to small, scattered foci (**Fig. 4A**). Occasional spatial proximity between SARS-CoV-2 and RV signals was observed, suggesting overlapping regions of susceptibility within the tissue.

**Figure 4.**
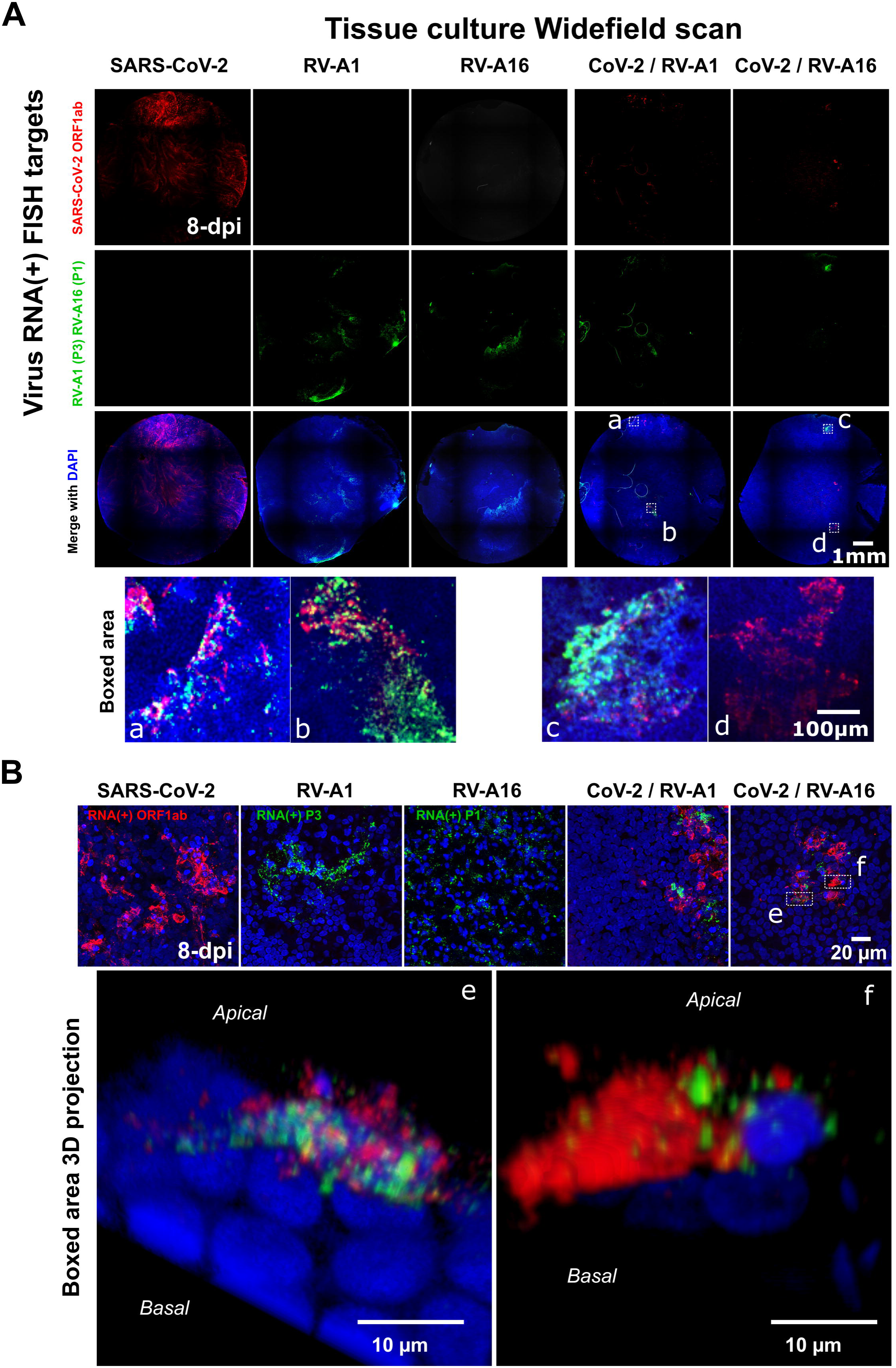
Spatial distribution of SARS-CoV-2 and RVs in coinfected tissue cultures. Single and coinfected bronchial epithelium were fixed at 8-days post infections and stained by RNA-FISH for SARS-CoV-2 RNA(+) ORF1ab, RV-A1 RNA(+) P3 region or A16 RNA(+) P1 region. Nuclei are stained with DAPI in blue. Tissue culture observation with widefield Olympus ScanR HCS microscope (**A**) and SP8 confocal microscope (Leica) (**B**).

Confocal microscopy was employed to investigate potential co-localization between SARS-CoV-2 and RVs. Intracellular RNA(+) puncta of SARS-CoV-2, RV-A1, and RV-A16 were predominantly localized near the apical epithelial surface. Analysis of overlapping regions revealed a small number of dual-infected cells; however, the absence of overlapping fluorescence signals indicates that viral RNA replication sites remain spatially segregated within coinfected cells (**Fig. 4B**). The majority of positive cells appeared to be infected by only one virus. Nevertheless, given the dense, multilayered architecture of the epithelium, which complicates precise cell segmentation, we cannot fully exclude the possibility that cells primarily infected by one virus may also harbor low levels of RNA(+) puncta from the other.

These findings suggest that coinfection constrains viral dissemination within the tissue, potentially through mechanisms that restrict productive infection of the same cell by both viruses.

### Strong inhibition of a superinfecting virus by a prior infection

To further explore mechanisms of cell exclusion, we performed staggered two-day superinfection experiments with RVs and SARS-CoV-2, a time point corresponding to peak replication for both viruses (**Fig. 1**). As a first step, we examined the distribution of RV-A1, RV-A16, and SARS-CoV-2 in tissue cultures single infections at 2 dpi, prior to superinfection. Viral RNA was detected by FISH staining and visualized with low-resolution, full-specimen scans. All viruses established as expected robust infections but showed distinct spread patterns. SARS-CoV-2–infected cells were widespread across the tissue, whereas RVs formed denser clusters (**Fig. S4**).

Next, we infected human bronchial epithelium cultures with SARS-CoV-2 (BA.2) or mock-infected them for 2 days, followed by superinfection with either RV-A1 or RV-A16 (**Fig. 5A**). Prior SARS-CoV-2 infection significantly impaired subsequent RV-A1 and RV-A16 superinfection, reducing RV titres and preventing long-term persistence, particularly for RV-A16 (**Fig. 5B and 5C**). In contrast to strict coinfection, SARS-CoV-2 replication was unaffected by superinfection with either RV-A1 or RV-A16.

**Figure 5.**
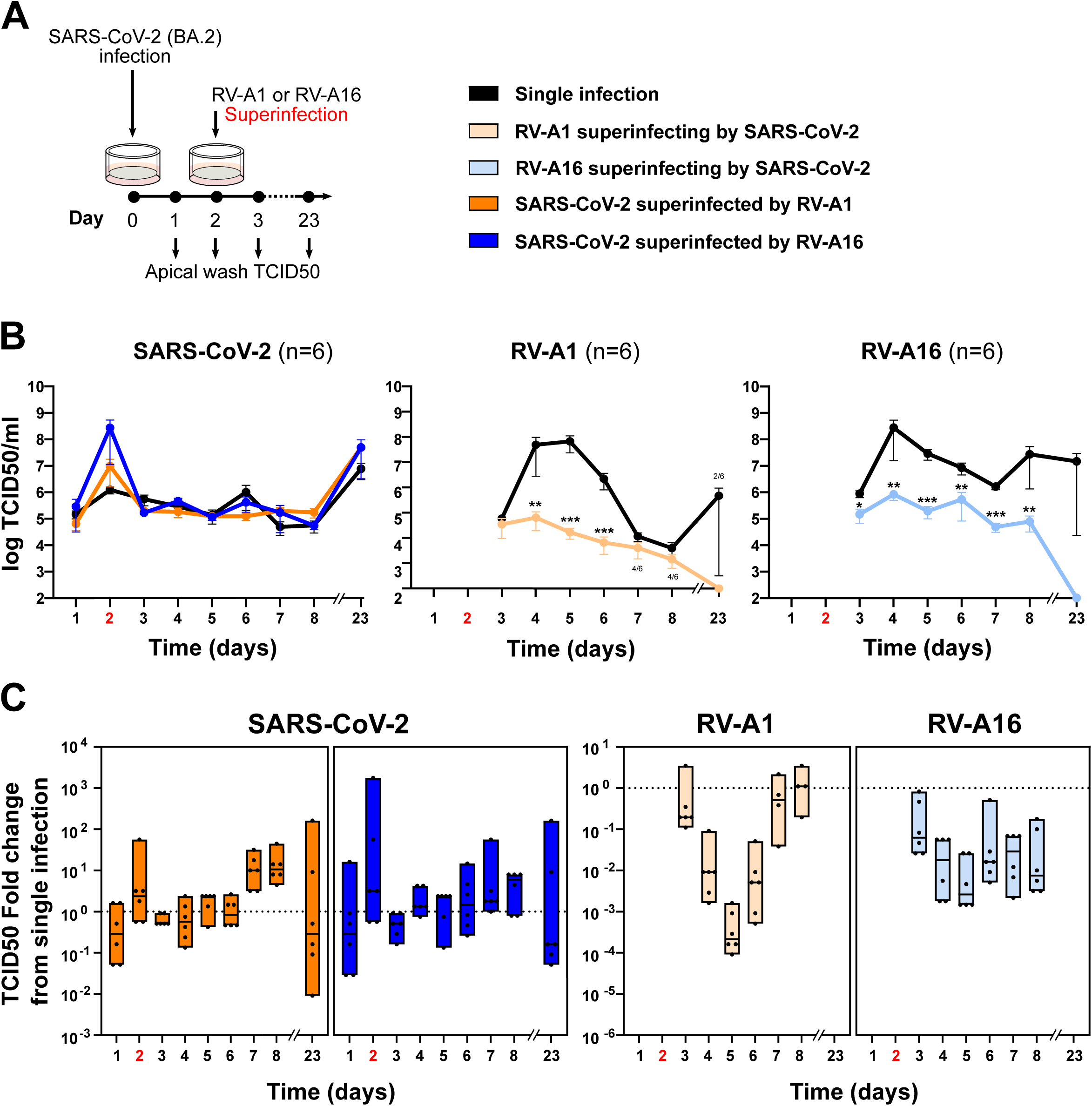
SARS-CoV2 (BA.2) infection interferes with a Rhinovirus superinfection. Primary bronchial epithelium tissue cultures were infected at day 0 by SARS-CoV-2 (BA.2) (10,000 TCID/tissue, 33.5°C) and then superinfected 2-days later by either RV-A1 or RV-A16 (10,000 TCID/tissue, 33.5°C). Single SARS-CoV-2 (BA.2) infection at 0-dpi, and single RV-A1 and RV-A16 infection at 2-dpi were done in parallel as control. Virus infectious progenies at the apical side have been differentially quantified at the indicated time points by TCID_50_ titrations. (**A**). RV-A1 and RV-A16 infectious progenies kinetics from single infection and superinfection are presented as mean±SEM of two independent experiments performed in three independent replicates each (**B**). Box-Whisker plots of the RV-A1 and RV-A16 titres fold changes between the single infected control (baseline) and the superinfection assay are represented (**C**). SARS-CoV-2 (BA.2) infectious progenies kinetics and change in titre (**D** and **E**). The number of samples with a measurable virus titre is indicated. Statistical comparison was performed using multiple t-test. *P* value for comparison across the groups are shown, *P < 0.05, **P < 0.01, ***P < 0.001. The 2-day timepoint, marking the superinfection, is highlighted in red. Apical wash collection at 2 dpi was performed prior to superinfection.

The reverse experiment, where cultures were first infected with RV-A1 or RV-A16 for 2 days before SARS-CoV-2 superinfection, showed the opposite outcome: strong interference against SARS-CoV-2 (**Fig. 6A**). This effect was stronger with RV-A16 than RV-A1 (**Fig. 6B and 6C**). SARS-CoV-2 superinfection of RV-A1 infected tissues persisted for all replicates, reaching by 23 days post-RV-A1 infection (21 days post-SARS-CoV-2 superinfection) comparable apical release to single-infection controls, consistent with RV-A1 non-persistence. At the contrary, few replicates of SARS-CoV-2 superinfecting of RV-A16 infected tissues persisted. However, persisting SARS-CoV-2 superinfection were associated with RV-A16 copersistence.

**Figure 6.**
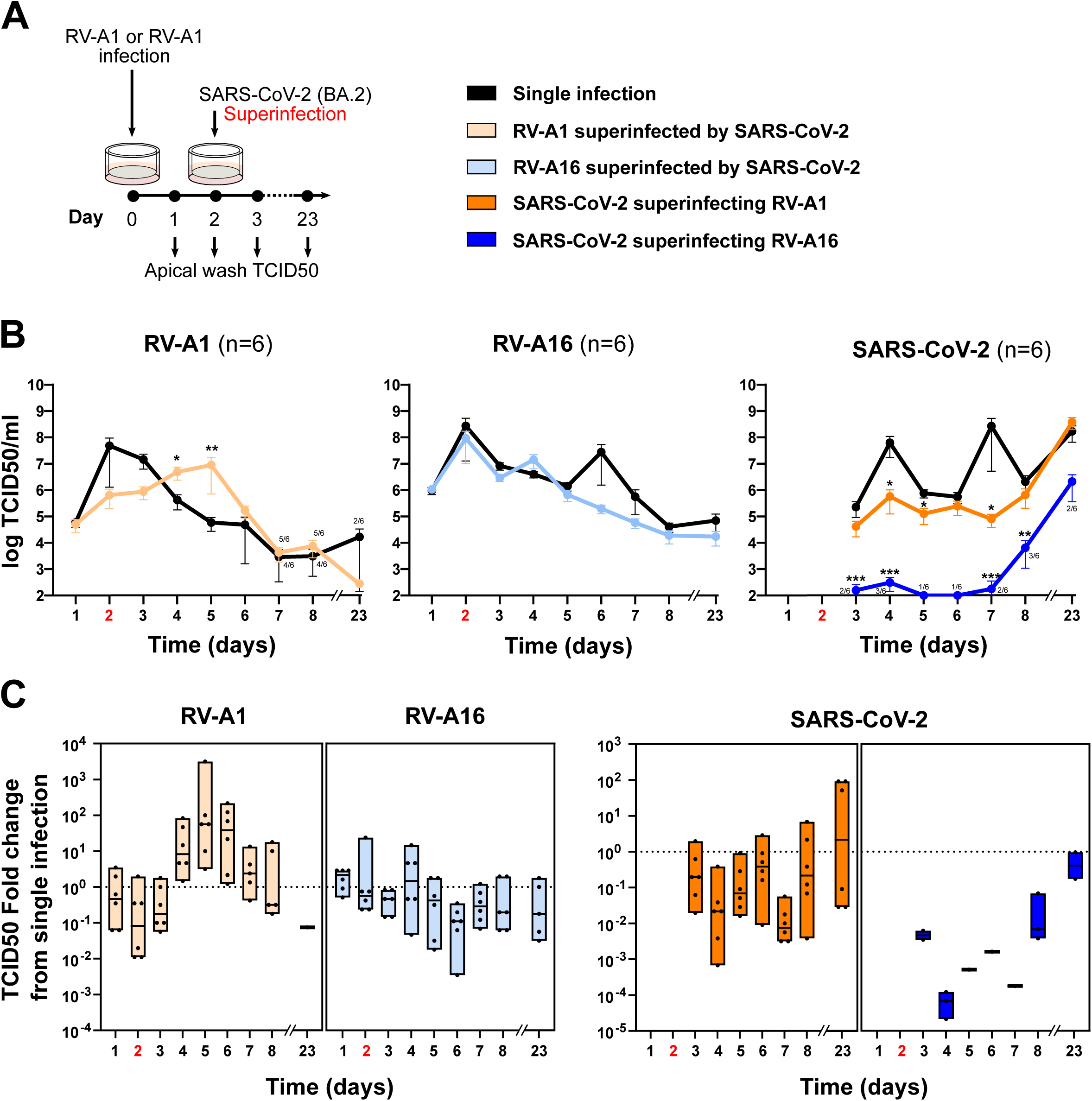
Rhinovirus pre infection interferes with the first steps of a SARS-CoV-2 (BA.2) superinfection. Primary bronchial epithelium tissue cultures were infected at day 0 by RV-A1 or RV-A16 (10,000 TCID/tissue, 33.5°C) and then superinfected 2-days later by SARS-CoV-2 (BA.2) (10,000 TCID/tissue, 33.5°C). Single RV-A1 and RV-A16 infection at 0-dpi, and single SARS-CoV-2 (BA.2) infection at 2-dpi were done in parallel as control. Virus infectious progenies at the apical side have been differentially quantified at the indicated time points by TCID_50_ titrations. (**A**). RV-A1 and RV-A16 infectious progenies kinetics from single infection and superinfection are presented as mean±SEM of two independent experiments performed in three independent replicates each (**B**). Box-Whisker plots of the RV-A1 and RV-A16 titres fold changes between the single infected control (baseline) and the superinfection assay are represented (**C**). SARS-CoV-2 (BA.2) infectious progenies kinetics and change in titre (**D** and **E**). The number of samples with a measurable virus titre is indicated. Statistical comparison was performed using multiple t-test. *P* value for comparison across the groups are shown, *P < 0.05, **P < 0.01, ***P < 0.001. The 2-day timepoint, marking the superinfection, is highlighted in red. Apical wash collection at 2 dpi was performed prior to superinfection.

Together, these results show that the extent of viral interference depends on both the timing of infection and the relative replication dynamics of the two coinfecting viruses.

### The inhibition of RV-A16 cell entry reduces interference against SARS-CoV2

To assess the impact of RV-induced superinfection exclusion on SARS-CoV-2, differentiated tissue cultures were apically inoculated with RV-A16 for 2 hours. Following this initial infection period, 1 µM Pleconaril, rhinovirus entry inhibitor, was applied to the basolateral medium to interfere with the dissemination of progeny RV-A16 virions, and renewed daily. At 2-days post RV-A16 infection, the tissue culture was superinfected by SARS-CoV-2 (BA.2) (**Fig. 7A**). A Pleconaril treatment control on SARS-CoV-2 done in parallel confirmed the absence of any antiviral effect of pleconaril against SARS-CoV-2 (**Fig. 7B**).

**Figure 7.**
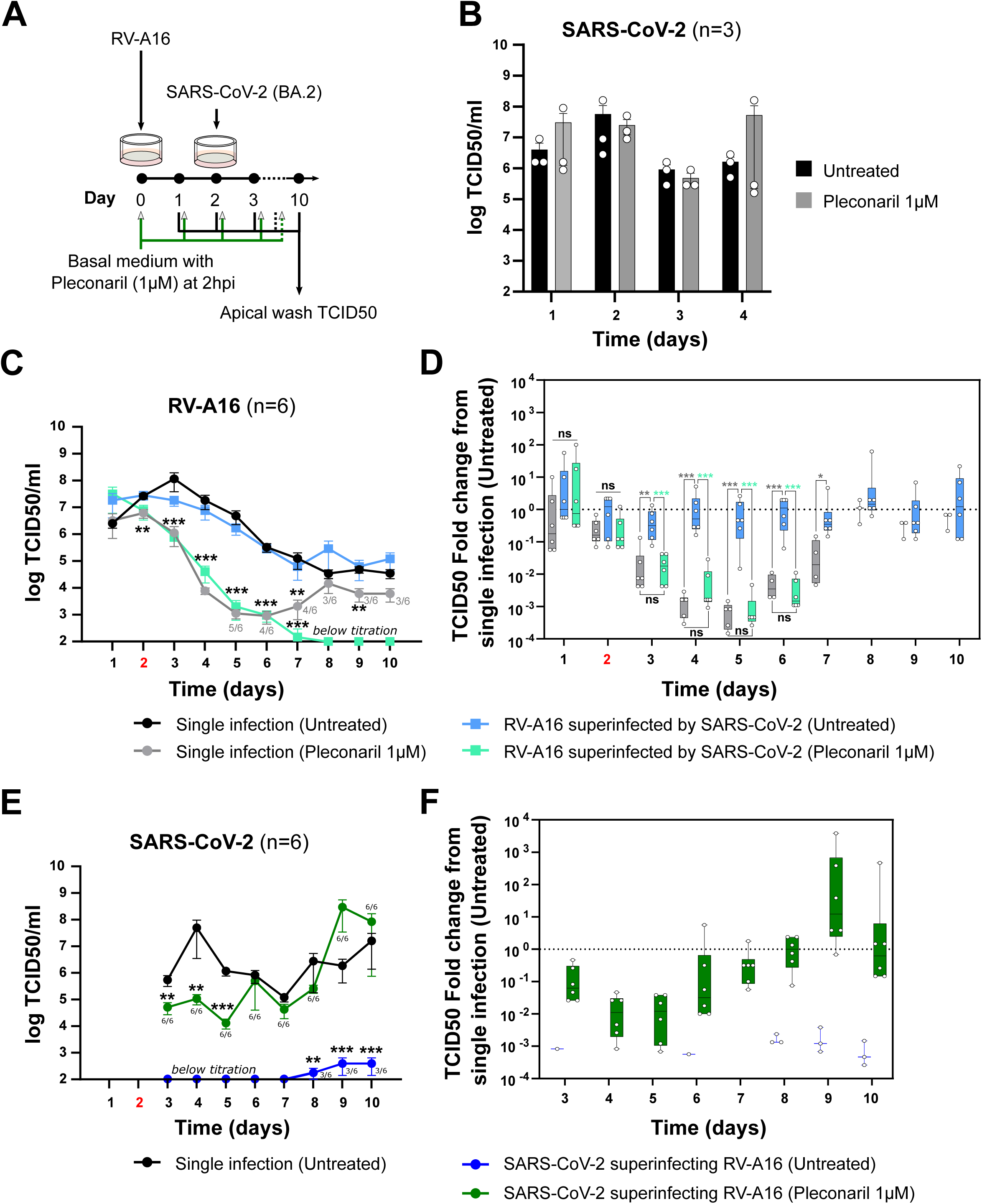
Specific inhibition of RV-A16 by Pleconaril potentiates SARS-CoV-2 (BA.2) superinfection. Primary bronchial epithelium tissue cultures were infected at day 0 by RV-A16 (10,000 TCID/tissue, 33.5°C), and treated with Pleconaril (1µM final) in the basolateral medium, starting at 2h post infection. Pleconaril was then administrated daily. At day-2, SARS-CoV-2 (BA.2) superinfection was performed (10,000 TCID/tissue, 33.5°C). Single RV-A16 and SARS-CoV-2 (BA.2) infections and mock treatment were done in parallel as control. Virus infectious progenies at the apical side have been differentially quantified at the indicated time points by TCID_50_ titrations. (**A**). SARS-CoV-2 (BA.2) single infection comparisons between Pleconaril treated and untreated are represented as mean±SEM of one tissue culture experiment performed in three independent replicate (**B**). Infectious progenies kinetics from single and superinfection assays and Box-Whisker plots of two independent experiments performed in three independent replicates each are represented for RV-A16 (**C** and **D**) and SARS-CoV-2 (**E** and **F**). Box-Whisker plots represent the corresponding virus titers fold changes between the mock treated single infection control (baseline) and the superinfection assay with or without Pleconaril treatment. The number of samples with a measurable virus titre is indicated. Statistical comparison was performed using multiple t-test. *P* value for comparison across the groups are shown, *P < 0.05, **P < 0.01, ***P < 0.001. The 2-day timepoint, marking the superinfection, is highlighted in red. Apical wash collection at 2 dpi was performed prior to superinfection.

Daily Pleconaril treatment exerted a strong antiviral effect against RV-A16, becoming significant at 2–3 days post-infection and resulting in up to a 3.5 log₁₀ reduction in viral titers (**Fig. 7C and 7D**). In the context of SARS-CoV-2 superinfection, Pleconaril treatment led to the complete clearance of RV-A16 by day 7, whereas approximately 50% of Pleconaril-treated RV-A16 single-infection replicates remained detectable up to 10 days post-infection. Consistently, untreated RV-A16 cultures superinfected with SARS-CoV-2 did not differ significantly from untreated RV-A16 single-infection controls (**Fig. 7C and 7D**).

In Pleconaril-treated cultures, SARS-CoV-2 superinfection was consistently established, producing higher viral titers than RV-A16 mock-treated controls between 3 and 5 days post-RV-A16 infection (equivalent to 1–3 days post-SARS-CoV-2 superinfection). Despite this enhancement, SARS-CoV-2 titers remained lower than those observed in the SARS-CoV-2 single-infection control. By day 6, when RV-A16 levels in Pleconaril-treated cultures had declined below the limit of quantification, no significant differences were observed between SARS-CoV-2 superinfection and SARS-CoV-2 single infection (**Fig. 7E and 7F**). In contrast, the persistence of RV-A16 in mock-treated controls continued to impose strong inhibition on SARS-CoV-2 replication with much lower viral titers and an established superinfection observed only in half of the replicates.

Finally, following co-infection settings, Pleconaril inhibition of RV-A16 tended to favor the co-infecting SARS-CoV-2; however, no significant differences were observed compared to the untreated co-infection control (**Fig. S5**). This could be attributed to the absence of a notable antiviral effect against RV-A16 prior 3-days of daily Pleconaril treatment (**Fig. 7C and 7D**). Taken together, our results indicate that the chemical inhibition of RV-A16 spread significantly favoured SARS-CoV-2 (BA.2) superinfection.

### Coinfections with non-structural protein chimeras of RV-A1 and RV-A16 do not show different levels of SARS-CoV-2 interference compared with the parental strains

Rhinovirus non-structural proteins are known to manipulate host cell structures and metabolism to enhance virus replication and propagation ^27,28,29^. A pairwise comparison of the non-structural protein sequences from the RV-A1 and A16 strains used in this study indicate an overall nucleotide identity of 78.84% and an amino acid identity of 87.86%. Nucleotide similarity variations range from 0.58 to 0.85 between RV-A1 and RV-A16 non-structural protein CDS (**Fig. S6**). To assess whether sequence variations in non-structural proteins between RV-A1 and RV-A16 may contribute to the stronger interference of RV-A16 against coinfecting SARS-CoV-2, we generated *in vitro* chimeric viruses by swapping the coding sequences of non-structural proteins between the RV-A1 and RV-A16 genomes. The resulting chimeric viruses were designated RV-A1-(A16) and RV-A16-(A1). The name in bracket indicates the genetic origin of the non-structural protein coding regions (**Fig. 8A**). These chimeras were then inoculated into bronchial epithelium tissue cultures, either as single infections or in coinfection with SARS-CoV-2 (BA.2).

**Figure 8.**
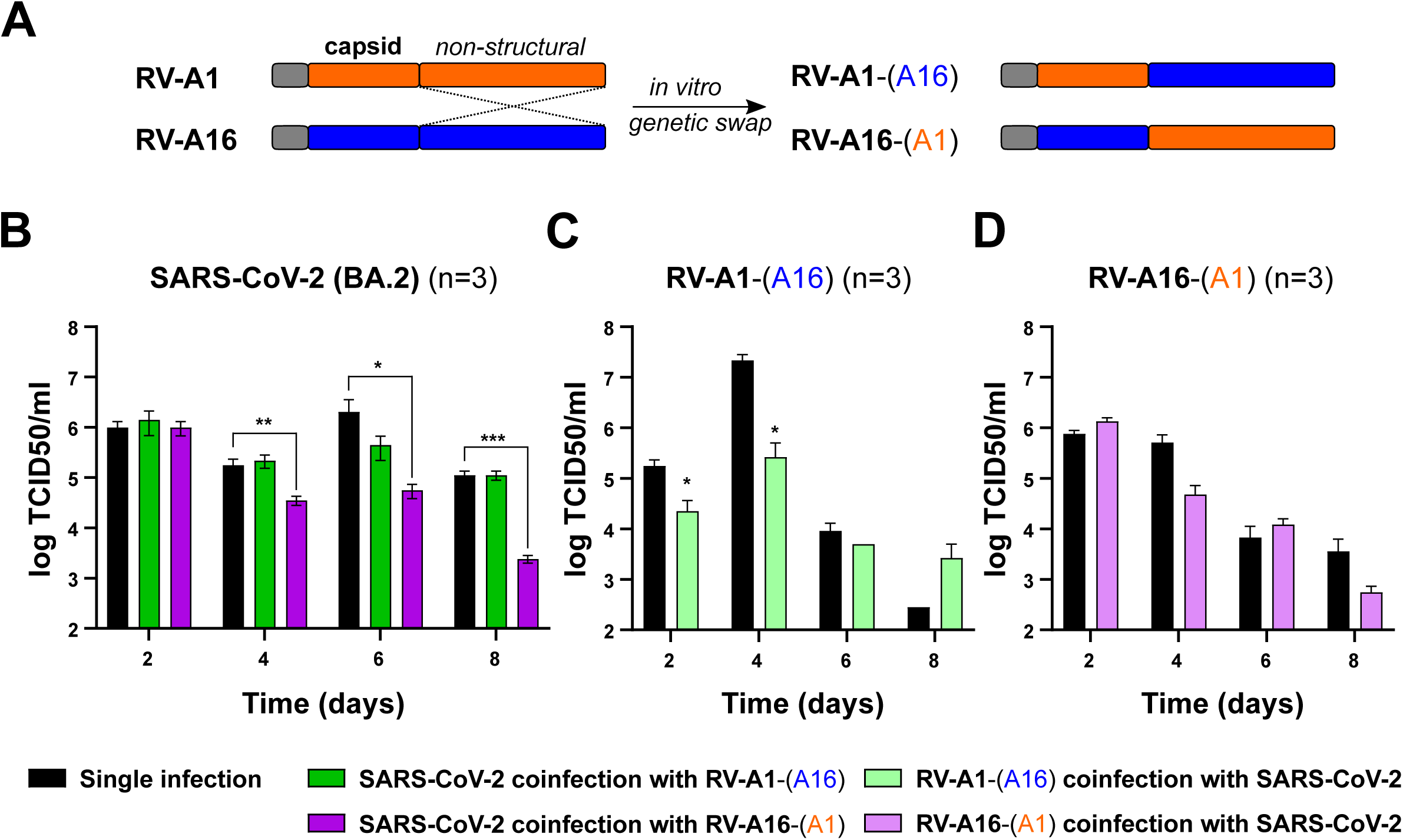
Rhinovirus non-structural proteins are not associated with the interference of a coinfecting SARS-CoV-2. The genomic coding sequences of the RV-A1 and RV-A16 non structural proteins were swapped *in vitro.* The resulting chimeric viruses were produced by transfection of *in vitro* transcribed swapped RNA in Hela Ohio Geneva cells. The rhinovirus chimeras were called RV-A1-(A16) and RV-A16-(A1), the name in bracket indicate the genetic origin of the non-structural proteins (**A**). Bronchial epithelium air-liquid cultures were single or coinfected by SARS-CoV-2 (BA.2), RV-A1-(A16), and RV-A16-(A1) at the apical surface (10,000 TCID50/tissue, 33.5°C). Virus infectious progenies at the apical surface were collected at the indicated time post infections and quantified by TCID_50_ titrations. Virus titres of the single infected control and the coinfected assay are represented as mean±SEM of one tissue culture experiment performed in three independent replicates each for SARS-CoV-2 (BA.2) (**B**), RV-A1-(A16) (**C**), and RV-A16-(A1) (**D**). Statistical comparison was performed using multiple t-test. *P* value for comparison across the groups are shown, *P < 0.05, **P < 0.01, ***P < 0.001.

Co-infection between the RV-A16-(A1) chimera and SARS-CoV-2 resulted in a significant reduction in SARS-CoV-2 infectious progenies at 4, 6, and 8 days post-infection (**Fig. 8B**). In contrast, co-infection between the RV-A1-(A16) chimera and SARS-CoV-2 did not show any significant differences (**Fig. 8B**). In parallel, SARS-CoV-2 did not induce particular interference against RV-A16-(A1) chimera, but significant reduction in RV-A1-(A16) chimera at 2- and 4-dpi (**Fig. 8C and 8D**). These results, in agreement with those obtained previously by using the WT RV-A1 and A16 progenitors, suggest that RV-A1 and A16 non-structural proteins are not associated with the magnitude of interference against coinfecting SARS-CoV-2.

## DISCUSSION

Our study provides evidence for viral interference between rhinoviruses (RVs), particularly RV-A1 and RV-A16, and SARS-CoV-2 Omicron (BA.2) in human primary bronchial tissue cultures. We found that the dynamics of co- and superinfections were shaped by both infection timing and replication levels, with RV-A16 exerting the strongest effects on SARS-CoV-2 replication at early stages. In contrast, SARS-CoV-2 had only modest effects on coinfecting RVs, although it demonstrated persistence in culture for up to 23 days, even in the presence of coinfecting RV. These findings were consistent across multiple donors, reinforcing the robustness of our observations. Our results extend prior work on respiratory viral coinfections, which has yielded conflicting outcomes. Some studies have reported protective effects of RVs (RV-A1 and –A16) against other respiratory viruses, including coronaviruses and influenza A virus (IAV) ^12,30,31^, whereas others described no measurable interference between RV-A16 and SARS-CoV-2 ^14^.

Notably, our findings differ from earlier reports examining pre-Omicron SARS-CoV-2 Wuhan or Alpha variants, which primarily relied on genome copy numbers as the measure of infection ^12,14^. While RT-qPCR offers sensitivity, speed, and lower labor requirements, it cannot distinguish between infectious and non-infectious particles, nor between complete and partial RNA genomes. This limitation is particularly relevant for coronaviruses, which generate multiple subgenomic RNAs through discontinuous transcription; as a result, RT-qPCR may underestimate interference effects, as we have previously reported with some antiviral compounds ^16,32^. By contrast, titration of infectious progeny provides a more direct and biologically relevant measure of viral spread and transmission potential. Variant-specific differences may further explain discrepancies across studies. Our profiling of innate immune factors revealed that coinfections prolonged secretion of type I/III interferons and cytokines such as IL-10, IL-22, IL-32, and TNFSF14. However, functional assays demonstrated that SARS-CoV-2 Omicron BA.2 replication was largely resistant to RV-A16 conditioned environment and to interferon, in contrast to ancestral Wuhan strains. This is consistent with recent evidence showing that Omicron variants have evolved enhanced interferon resistance relative to early pandemic isolates ^33,34^. RV-A16 elicited an earlier and stronger innate immune response than RV-A1. Furthermore, RV-A1 was highly sensitive to interferon, in contrast to RV-A16, highlighting intertypic differences among rhinoviruses. Together, these results emphasize the importance of investigating contemporary SARS-CoV-2 variants, which may engage differently with common respiratory viruses than their predecessors, and suggest that interferon responses during coinfection are more likely a consequence of interference rather than its primary cause. In the specific case of RV-A16 and SARS-CoV-2 coinfection, the robust IFN response induced by RV-A16 may be prolonged in time, as RV-A16– mediated interference would likely delay the onset of SARS-CoV-2 counter IFN measures.

Our RNA FISH experiments revealed distinct but adjacent viral foci, with only a limited presence of dual-infected cells, strongly suggesting that cell superinfection exclusion is a central mechanism underlying viral interference. Similar phenomena have been reported for progeny viruses in single infections of influenza, C-hepatitis and coronaviruses ^35,36,37^.Viruses may employ multiple strategies to achieve superinfection exclusion, including early-stage mechanisms that prevent further virus adsorption and later-stage mechanisms that interfere with viral genome translation or replication ^38,39^. In the context of SARS-CoV-2 and rhinovirus coinfections, our findings indicate that established infection by either virus restricts subsequent superinfection, consistent with these mechanisms. This result aligns with recent clinical observations analyzing 10,493 nasal swabs from 1,156 participants in the Human Epidemiology and Response and showing that pre-existing rhinovirus infection reduces susceptibility to SARS-CoV-2 ^40^. However, our data also suggest the reverse scenario, where pre-existing SARS-CoV-2 infection can limit rhinovirus superinfection.

Inhibition of RV-A16 entry and spread *via* Pleconaril reduced its interference with SARS-CoV-2, further supporting the role of viral entry and dissemination dynamics in shaping the observed interference. While at the contrary genetic swaps of the non-structural proteins coding regions between RV-A1 and A16 does not change the magnitude of the observed interference. Nevertheless, further investigations are necessary to determine the molecular mechanisms of interference between rhinovirus and SARS-CoV-2 and to decipher if it occurs at the level of viral entry, post-entry events, or a combination of both. Interestingly, despite an initial suppression by RVs, SARS-CoV-2 displayed a capacity for long-term persistence and eventual recovery of replication to levels comparable with single-infection controls. This resilience may plausibly reflect the ability of SARS-CoV-2 to exploit preserved susceptible cell populations, underscoring the complexity of virus–virus interactions in which suppression may be transient rather than permanent. However, the absence of immune cell populations limits the extent to which our findings can be directly extrapolated to *in vivo* settings, where adaptive and innate immune responses are likely to influence both viral persistence and the outcome of coinfections. Nevertheless, the observed resilience of SARS-CoV-2 to rhinovirus coinfection aligns with epidemiological observations. Both RVs and SARS-CoV-2 are detected year-round, yet often display inverse prevalence trends, supporting the existence of viral interference at the population level ^40,41,42,43^.

Another observation, based on microscopy and absence of leakage, was the maintenance of epithelial tissue integrity across all infection settings. The preservation of structural integrity even during peak replication suggests a high degree of epithelial resilience ^44,45^. The regenerative capacity of lung epithelium is ensured by the presence of a resident population of basal stem cells able to self-renew following injuries and to differentiate into the cell types constituting the epithelium ^44,45,46^. A limitation of our work is that we did not determine whether infected tissue maintenance was complete or only limited to cell self-renewal without proper cell differentiations. At least, the sustained presence of SARS-CoV-2 suggests the preservation of ciliated cells, which are known to express high levels of SARS-CoV-2 receptor ACE2, both *in vitro* and in human airway biopsies ^47^. Clarifying the nature and the mechanisms of tissue maintenance during persistent or recurrent infections is clinically relevant. Dysregulation of lung epithelium regenerative process could have implications for chronic respiratory conditions such as COPD or asthma, which are known to be exacerbated by RV infections ^2,5^, or some observed long-COVID lung tissue sequelae ^48^.

Taken together, our findings suggest that interference between RVs and SARS-CoV-2 is a multifaceted phenomenon driven by viral load, type, timing, and multiple exclusion mechanisms. It could be hypothesized that viral interference may act as an infectious bottleneck, which could impact viral genetic diversity and lead to new evolutionary trajectories in the context of multi-infection compared to a single infection context. Common respiratory viruses, like RVs could have participated to shape the epidemiology of SARS-CoV-2 variants by altering their replication dynamics within the respiratory tract. Care must be taken in that hypothesis as our study was performed in *ex vivo* bronchial tissue cultures, which, while physiologically relevant, may not fully capture the complexity of *in vivo* coinfections. Additionally, the precise molecular determinants of interference remain to be defined. Future work should focus on dissecting these mechanisms at the single-cell level, including the role of receptor availability, cellular antiviral states, and competitive use of host resources.

## METHODS

### Cell lines culture

Human cervix adenocarcima HeLa-Ohio Geneva (HOG) (ECACC 84121901) cells were obtained from Pr. Laurent Kaiser (University Hospital Geneva, Switzerland). Monkey kidney epithelial VeroE6 cells transfected with transmembrane serine protease 2 (TMPRSS2) were obtained from Pr. Volker Thiel (University of Bern, Switzerland). Cells were cultured in Dulbecco’s modified Eagle’s medium (DMEM; cat#D6429; Sigma-Aldrich St. Louis, USA) supplemented with 10% fetal bovine serum (FBS; cat#10270; Invitrogen, Carlsbad, USA) and 1% non-essential amino acids (NEAA; cat#M7145; Sigma-Aldrich), washed in PBS, and detached with trypsin-EDTA (cat#C-41020; Sigma-Aldrich). Cell lines were cultured at 37°C under a humid atmosphere containing 5% CO_2_.

### Human *in vitro* differentiated bronchial epithelia tissue cultures at air-liquid interface

The human bronchial explants were obtained from individual donors (Donor 793, 839, and 859; Epithelix SA, Geneva, Switzerland). Experiments were primarily conducted using Donor 793 bronchial explants. Bronchial explants primary cells have been seeded on Type IV collagen (cat#C5533; Sigma Aldrich) coated microporous permeable membrane (0.4 µm pore) of Transwell ® device (24-well plate; 3470; Costar Corning New York, USA) and first expanded at liquid-liquid interface with PneumaCult™-Ex Basal Medium (cat#05009; Stemcell; Basel Switzerland) supplemented with 1X PneumaCult™-Ex 50X-Supplement (cat#05019; Stemcell) and 0.2X hydrocortisone (cat#07925; Stemcell). Then cells were differentiated for four weeks at air-liquid interface (ALI) with maintenance medium composed of PneumaCult™-ALI Base Medium (cat#05002; Stemcell,) supplemented with 1X of PneumaCult™-ALI 10X supplement (cat#05003; Stemcell), 1X hydrocortisone (cat#07925; Stemcell), 4µg/mL heparin (cat#07980; Stemcell), and 150 ng/mL retinoic acid (cat#R2625; Sigma Aldrich) in the basolateral compartment. Cells in transwell ® were cultivated at 37°C under a humid atmosphere containing 5% CO2.

### Viruses

The SARS-CoV-2 Omicron (BA.2) variant was obtained from the Robert Koch Institute (Berlin, Germany) through European Virus Archive global (EVAg). SARS-CoV-2 ―Wuhan‖ (TAR clone 3.3, München-1.1/2020/929) were obtained from Dr. Volker Thiel (University of Bern) ^49^. The RV-A1 and RV-A16 were a gift from Dr. Wai Ming Lee (Department of Pediatrics, School of Medicine and Public Health, University of Wisconsin-Madison). SARS-CoV-2 was expanded at 37°C on VeroE6/TMPRSS2 cells and rhinoviruses were expanded at 33.5°C on HOG cells, all were in DMEM medium supplemented with 4% of FBS, and 1X nonessential amino acids. Virus titers were determined on their respective cell lines by calculating the 50% tissue culture infective dose (TCID50) according to the Spearman-Kärber method. All experiments were carried out in a biosafety level-3 laboratory.

### Virus infection

Infection of differentiated bronchial epithelium tissues cultures were carried out as previously described^32^. Prior to infection, a 20-minutes apical wash was performed by adding 200 μl of ALI maintenance medium to the apical side. Inserts were inoculated on the apical side with 10,000 TCID50 of the indicated virus in a final volume of 100 μl for 2-hours at 33.5°C under a humid atmosphere containing 5% CO2. For coinfection, both viruses were mixed in the same inoculum with 10,000 TCID50 each in a final volume of 100 μl. After incubation, virus inocula were removed, apical surfaces quickly washed two-times with PBS and tissue cultures incubated at 33.5°C under a humid atmosphere containing 5% CO2. Virus produced at the apical side was collected at different time points post infection by 20 min apical wash with 200 µl of ALI maintenance medium at 33.5°C. Collected viruses were quantified by TCID50 titration with Vero E6/TMPRSS2 cells for SARS-CoV-2, and HOG cells for RV-A1 and A16. The basolateral medium was changed after each apical sample collection. All collected samples were conserved at −80°C until use. All experiments were carried out in a biosafety level-3 laboratory. Interferon (IFN) and pleconaril treatments were done at the indicated times pre and/or post-infection for the indicated durations and at the indicated concentrations in fresh basolateral medium. The human IFN-α2a was obtained from PBL Assay Science (cat#11100), the human IFN-λ1 and λ2 was obtained from PeproTech (cat#300-02L and −02K; Thermo Fisher; Waltham, USA). Pleconaril was obtained from Sigma Aldrich (SML0307) and dissolved in DMSO.

### RNA FISH with branched DNA signal amplification

Viral RNAs were stained by fluorescence *in situ* hybridization (FISH) as previously described ^16^. Following 30 min fixation with 4% PFA in PBS at RT and three washes in PBS, cells were permeabilized with absolute ice-cold methanol overnight at −20°C. Samples were then rehydrated by incubation with 75%, 50%, 25%, 0% of ice-cold methanol and PBST (PBS supplemented with 0.1% of Tween-20) for 5 min each, then washed on ice with a vol/vol 5XSSCT/PBST solution for 5min, and afterwards with a 5XSSCT solution for 5min. Virus RNAs were then FISH-stained using ViewRNA mRNA FISH assay according to the manufacturer’s instructions (Thermo Fisher). Custom-made virus FISH probes were directed against the ORF1a sequences between positions 401-1327 for SARS-CoV-2, and against the P1 sequences for RV-A1 and RV-A16 (Thermo Fisher). After staining, microporous membranes were detached from their transwell support and mounted between slide/coverslip with ProLong mounting media (P10144; ThermoFisher). Cells, facing the coverslip side, were imaged using a SP8 confocal microscope (Leica) and Widefield - Olympus ScanR HCS microscope.

### Rhinovirus chimeric genomes engineering

Chimeric genomes were engineered with a previously described fusion-PCR-based procedure ^50,51^. The RV-A1 and RV-A16 RNAs were extracted with the Virus RNA Min Elute extraction kit (cat#11858882001; Roche, Mannheim, Germany), denatured (5 min at 65°C), and then subjected to full-length reverse transcription (30 min at 50°C) with a Maxima H Minus First Strand cDNA synthesis kit (cat#K1652; Thermo Fisher) with specific 3’-end reverse primers. Full-length cDNA were used as template for producing overlapping segments by nested PCR with a Phire Hot Start Pol II kit (cat#F122S; Thermo Fisher). The amplified overlapping segments were subjected to electrophoresis in an agarose gel, purified with a Relia Prep^TM^ DNA Clean-Up and Concentration system (cat#A2893; Promega; Madison, USA), and quantified with a NanoDrop spectrophotometer (Thermo Fisher). Equimolar amounts of purified overlapping segments were ligated without primers with the Phire Hot Start Pol II. The ligated chimeric genomes were then amplified in the same reaction mixture by adding specific T7-promoter tailed 5’-end primer and poly(T) tailed 3’end primers. The amplified T7 promoter/chimeric rhinovirus genomes were purified with a Wizard ® SV Gel and PCR Clean-Up System (Promega) and *in vitro* transcribed (2-hours at 37°C) with the HiScribe™ T7 High Yield RNA Synthesis Kit (cat# E2040S; NEB; Ipswich, USA). Generated T7-promoter chimeric rhinovirus RNAs were subjected to DNAse I treatment for 15 min at 37°C (Roche), and purified with the quick RNA columns (cat#R1058; Zymo research; Irvine, USA). Hela Ohio Geneva (HOG) cells were transfected with 1 μg of purified chimeric rhinovirus RNAs in OptiMEM medium (cat#11058-021; Thermo Fisher) by using Lipofectamine-RNAi Max transfection reagent (cat#1153257; Invitrogen). Transfected cells were incubated at 33.5°C until complete cell lysis. Cell supernatants were collected, clarified by centrifugation, and passaged once on 80% confluent HOG cells grown in T-75 flasks incubated at 33.5°C. A list of primers is detailed in **supplementary table S1**.

### Inflammation factors multiplex microbead-based immunoassay

Collected basolateral supernatants from infected human bronchial epithelium tissue cultures were inactivated by 0.5% β-propiolactone (cat# B23197; Alfa Aesar; Haverhill, USA) and incubated overnight at +4°C. Following sample inactivation, β-propiolactone was hydrolysed (2-hours at 37°C), inflammation factor levels were analyzed with the Bio-Plex ProTM Inflammation Panel 1, 37-Plex (cat#171-AL001M, Standard lot#64530752, Control lot#64593524; Bio-Rad, Cressiet, Switzerland) in 50µl of basolateral supernatants, according to manufacturer’s instructions. The lecture of the results was performed with Bio-Plex® 200 Multiplex system (Bio-Rad).

### Statistical methods

Statistical comparisons across groups were carried out by the Student’s t-test. A *p-value* < 0.05 was considered statistically significant. Statistical analyses were performed using the GraphPad Prism software (Version 10.4.2).

## DATA AVAILABILITY

The data that support the findings of this study are available from the corresponding author upon reasonable request.

## ACKNOWLEDGMENTS

We gratefully acknowledge Prof. Urs F. Greber for financial support through the Swiss National Science Foundation (SNSF Covid Special Call Grant No. 31CA30_196177/1) and the Coronavirus Research Grant from the University of Zurich. We also thank Prof. Urs F. Greber, Prof. Cornel Fraefel, and Dr. Maarit Suomalainen for their critical review of the manuscript and valuable advice. Finally, we acknowledge Sara Costa Fernandes for preparing the human primary bronchial tissue cultures.

## AUTHOR CONTRIBUTION

R.V. conceived the project, designed and performed the experiments, analysed data, and wrote the manuscript.

