## Supplemental Table 1 for "DYNAMICS OF RHINOVIRUS SARS-COV-2 COINFECTIONS AND SUPERINFECTIONS IN HUMAN AIRWAY CULTURES REVEAL TYPE-DEPENDENT VIRAL INTERFERENCE"

**Supplementary table 1**

| **Name** | **Sequence 5’-3’** | **Position ^a^** |
| --- | --- | --- |
| **5’ and 3’ ends genomes primers** |  |  |
| RVA1-T7prom-1F | AAAGGGAAGCTTTAATACGACTCACTATAGGG**TTAAAACTGGGTGTGGGTTGTTCCCA** | 5’end |
| RVA1-end-R | **(T)_24_** **ATAGAATTAAAGAATCATTCATTCATTATTTCTATATCTAAAATTTTTCATACC** | 3’end |
| RVA16-T7prom-1F | AAAGGGAAGCTTTAATACGACTCACTATAGGG**TTAAAACTGGATCTGGGTTGTTCCCA** | 5’end |
| RVA16-end-R | **(T)_24_** **ATAAAACTAACAAACTATTCATTTATCAATTCTATATCTAGAATTTTTCATACCAC** | 3’end |
| **RV-A1-(A16) overlapping primers** |  |  |
| RVA1(r)3197R | *AGATTTCTGTATATTAGATTACCAACATGCACATACATGTCACTAGGCCC***AGCAGTTGTTATAGTGTTTCTGCGG** | 3173-3197 |
| RVA16(r)3185F | *CAGGTGACGTGACAACAGCCATAGTCCGCAGAAACACTATAACAACTGCT***GGGCCTAGTGACATGTATGTGCATG** | 3185-3209 |
| **RV-A16-(A1) overlapping primers** |  |  |
| RVA16(r)3185R | *AAGTTTCTATATATTAAGTTACCTACATGCACATATAGATCACTGGGCCC***AACAGTTGTTAGATTTGTTCTAGGT** | 3160-3184 |
| RVA1(r)3197F | *AAGTACACAATGATGTGGCTATAAGACCTAGAACAAATCTAACAACTGTT***GGGCCCAGTGATCTATATGTGCATG** | 3198-3222 |

**^a^** Positions are according to the acceptor genome

T7 promotor sequence is in underlined

Priming nucleotides with the acceptor genome are in **bold**

Non-priming 5’ tails of the donor genome located upstream or downstream the 1D/2A breakpoint are in *Italic*
