## Supplemental Figure 1 for "DYNAMICS OF RHINOVIRUS SARS-COV-2 COINFECTIONS AND SUPERINFECTIONS IN HUMAN AIRWAY CULTURES REVEAL TYPE-DEPENDENT VIRAL INTERFERENCE"

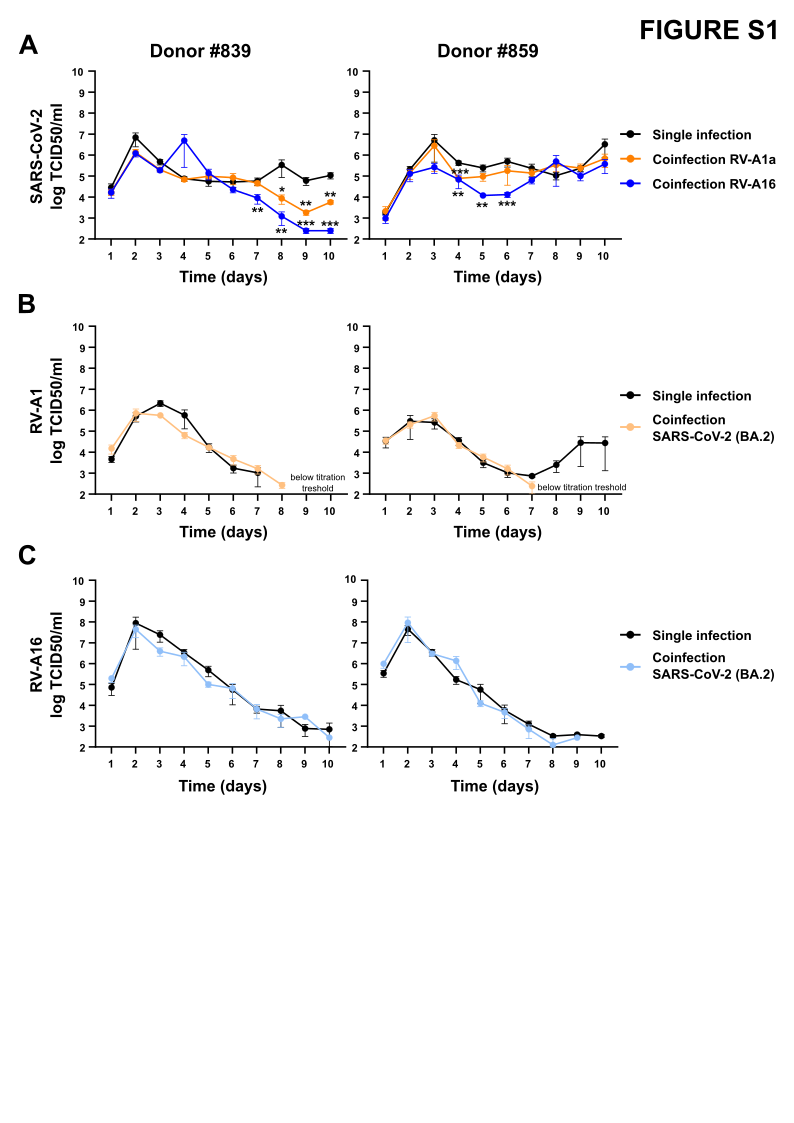

**Figure S1. SARS-CoV-2 and Rhinoviruses coinfections of primary bronchial epithelium tissue culture from different donors**. Two different single donor bronchial epithelium air-liquid cultures (donor#839 and #859) were single or coinfected by SARS-CoV-2 (BA.2), RV-A1, and RV-A16 at the apical surface (10,000 TCID_50_/tissue, 33.5°C). Virus infectious progenies at the apical surface were collected by apical wash at the indicated time post infections. Apical infectious virus progenies were differentially quantified by TCID_50_ titrations on VeroE6-TMPRSS (SARS-CoV-2) and Hela Ohio Geneva (Rhinoviruses) cells. Single and coinfection kinetics are presented for each virus as mean±SEM of two independent experiments performed in three independent replicates each. Statistical comparison was performed using multiple t-test. *P* value for comparison across the groups are shown, *P < 0.05, **P < 0.01, ***P < 0.001.
