## Supplemental Figure 2 for "DYNAMICS OF RHINOVIRUS SARS-COV-2 COINFECTIONS AND SUPERINFECTIONS IN HUMAN AIRWAY CULTURES REVEAL TYPE-DEPENDENT VIRAL INTERFERENCE"

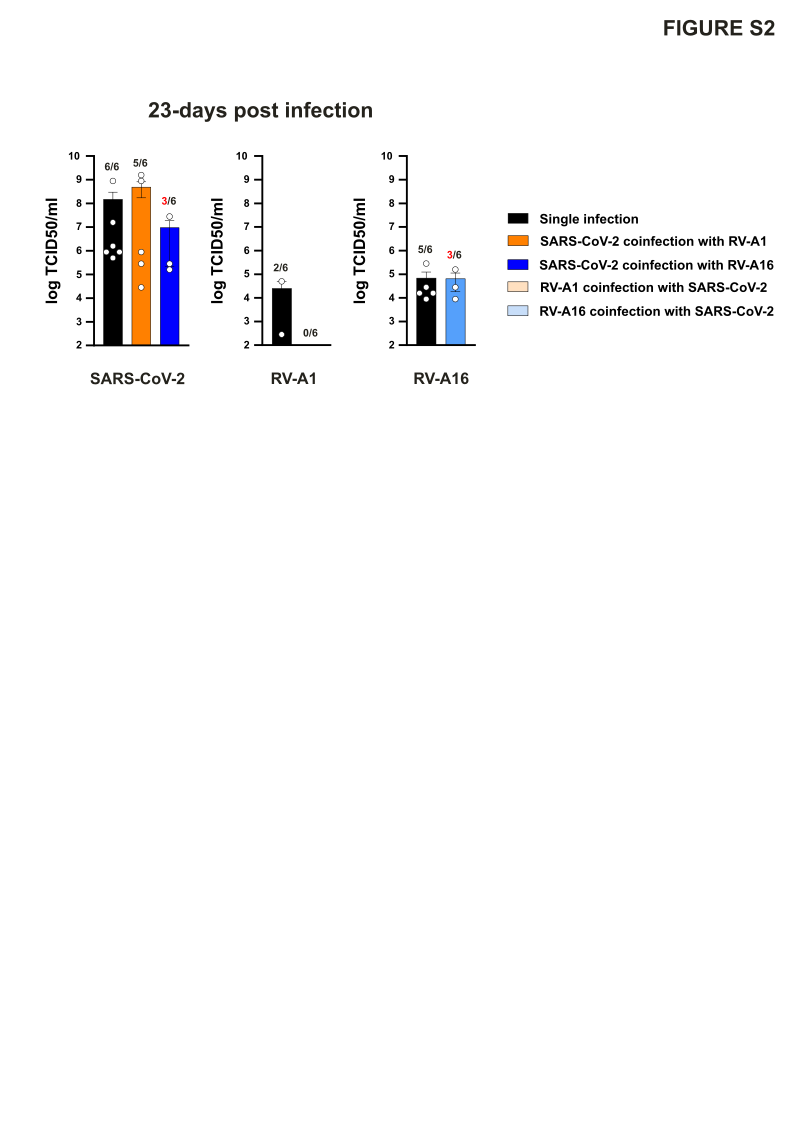

**Figure S2. SARS-CoV-2 and Rhinoviruses long-term coinfections of primary bronchial epithelium tissue culture.** Single donor bronchial epithelium air-liquid cultures were single or coinfected by SARS-CoV-2 (BA.2), RV-A1, and RV-A16 at the apical surface (10,000 TCID_50_/tissue, 33.5°C). Apical infectious virus progenies at 23-days post infection were differentially quantified by TCID_50_ titrations on VeroE6-TMPRSS (SARS-CoV-2) and Hela Ohio Geneva (Rhinoviruses) cells. Mean±SEM of two independent experiments performed in three independent replicates each. The number of samples with a measurable virus titre is indicated. For coinfections involving SARS-CoV-2 and RV-A16, samples with detectable virus titers are highlighted in red to indicate that they originate from the same coinfected tissue cultures.
