## Supplemental Figure 3 for "DYNAMICS OF RHINOVIRUS SARS-COV-2 COINFECTIONS AND SUPERINFECTIONS IN HUMAN AIRWAY CULTURES REVEAL TYPE-DEPENDENT VIRAL INTERFERENCE"

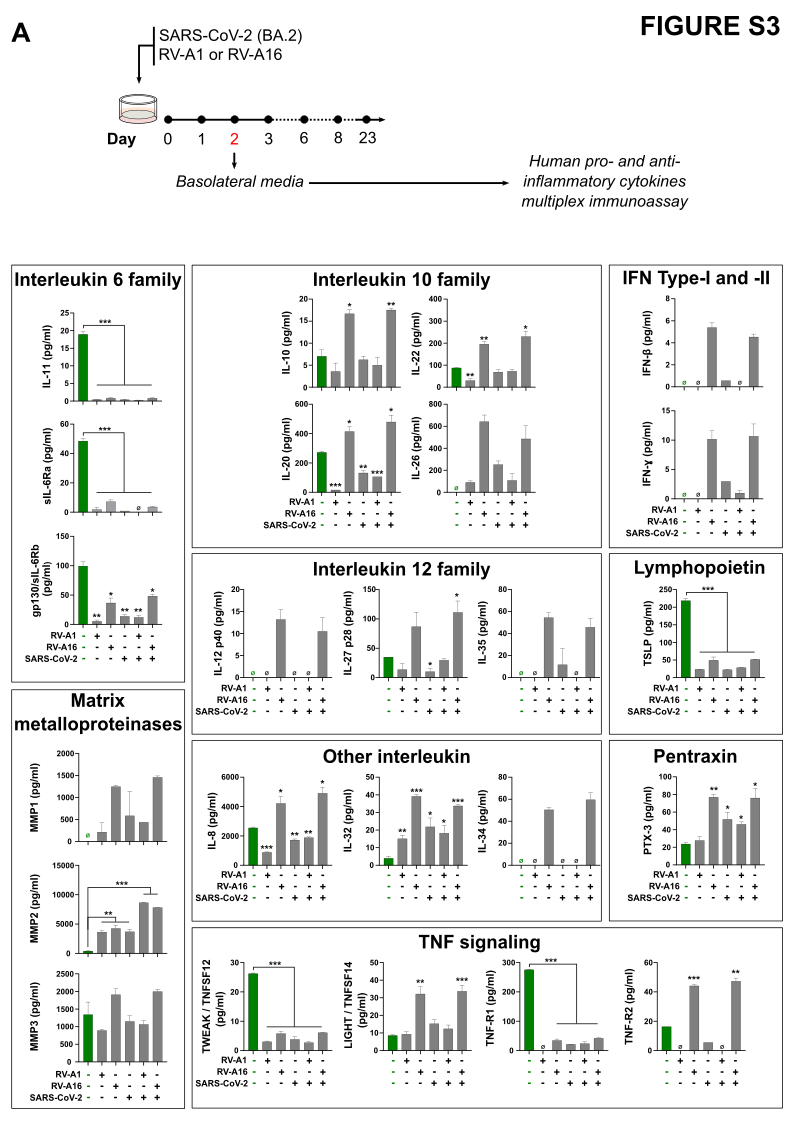

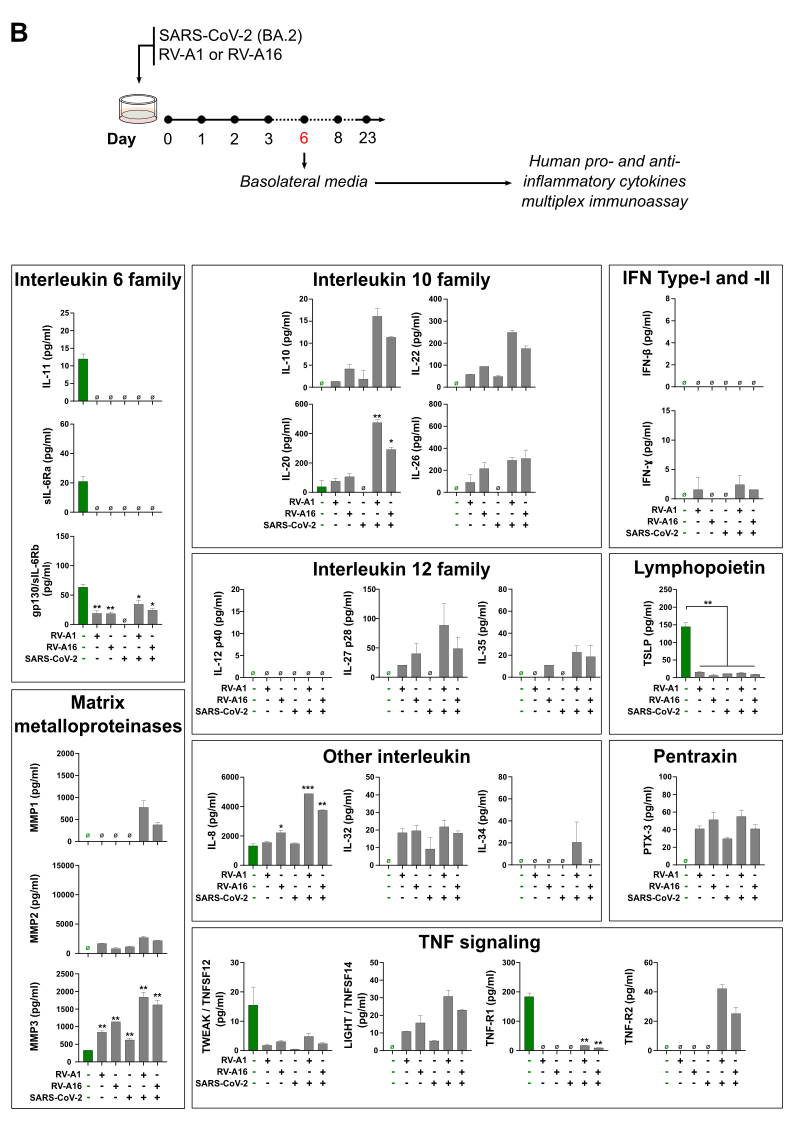

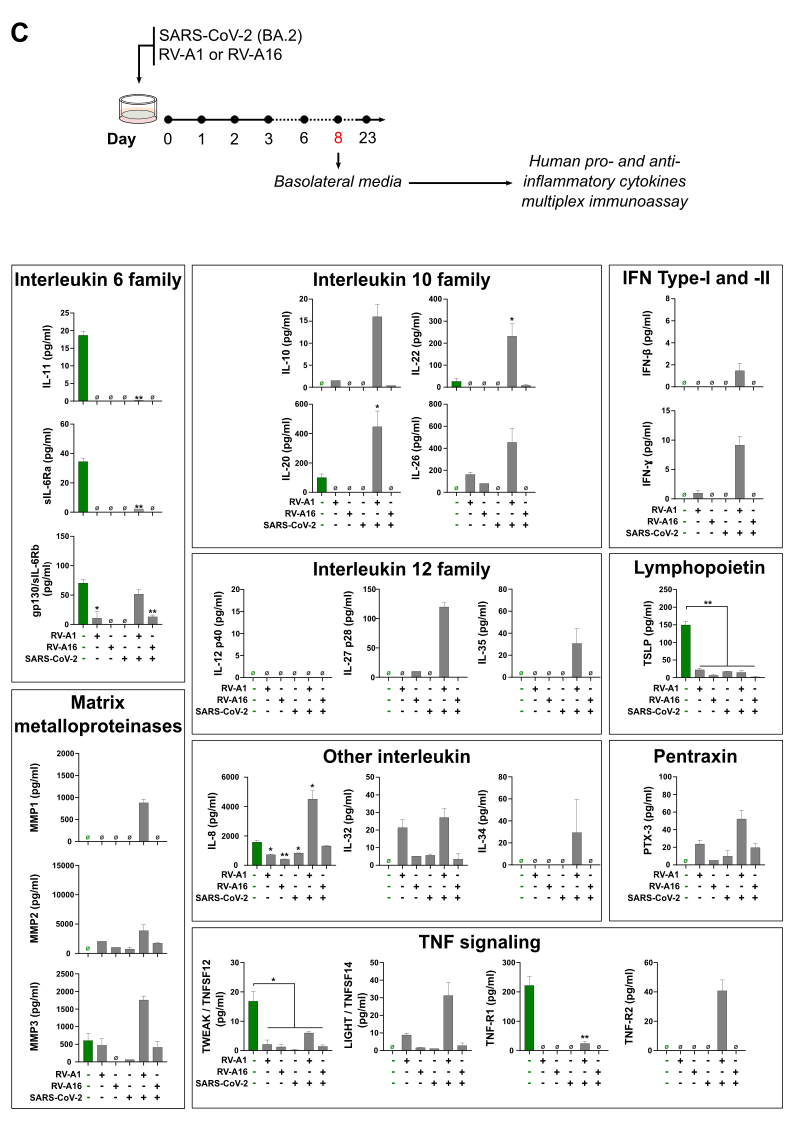

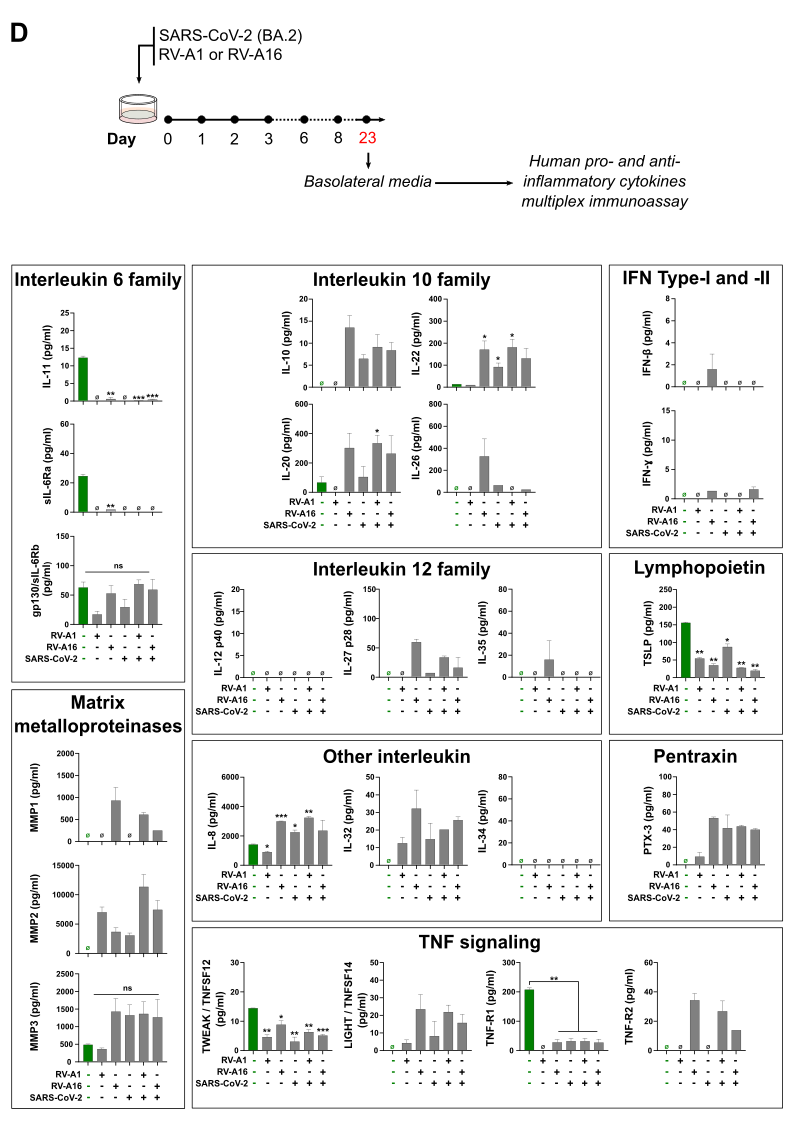
**Figure S3.** **Profiles of the innate immunity factors secreted by infected primary bronchial epithelium tissue culture.** Innate immune factor concentrations in basolateral media collected at 2- (**A**), 6- (**B**), 8- (**C**) and 23-days post infection (**D**) were captured by multiplex microbead-based immunoassay and analyzed. Data are the mean of two independent replicates. Statistical comparison was performed using multiple t-test. *P* value for comparison between the mock infected controls and the infected groups are shown, *P < 0.05, **P < 0.01, ***P < 0.001.
