## Supplemental Figure 4 for "DYNAMICS OF RHINOVIRUS SARS-COV-2 COINFECTIONS AND SUPERINFECTIONS IN HUMAN AIRWAY CULTURES REVEAL TYPE-DEPENDENT VIRAL INTERFERENCE"

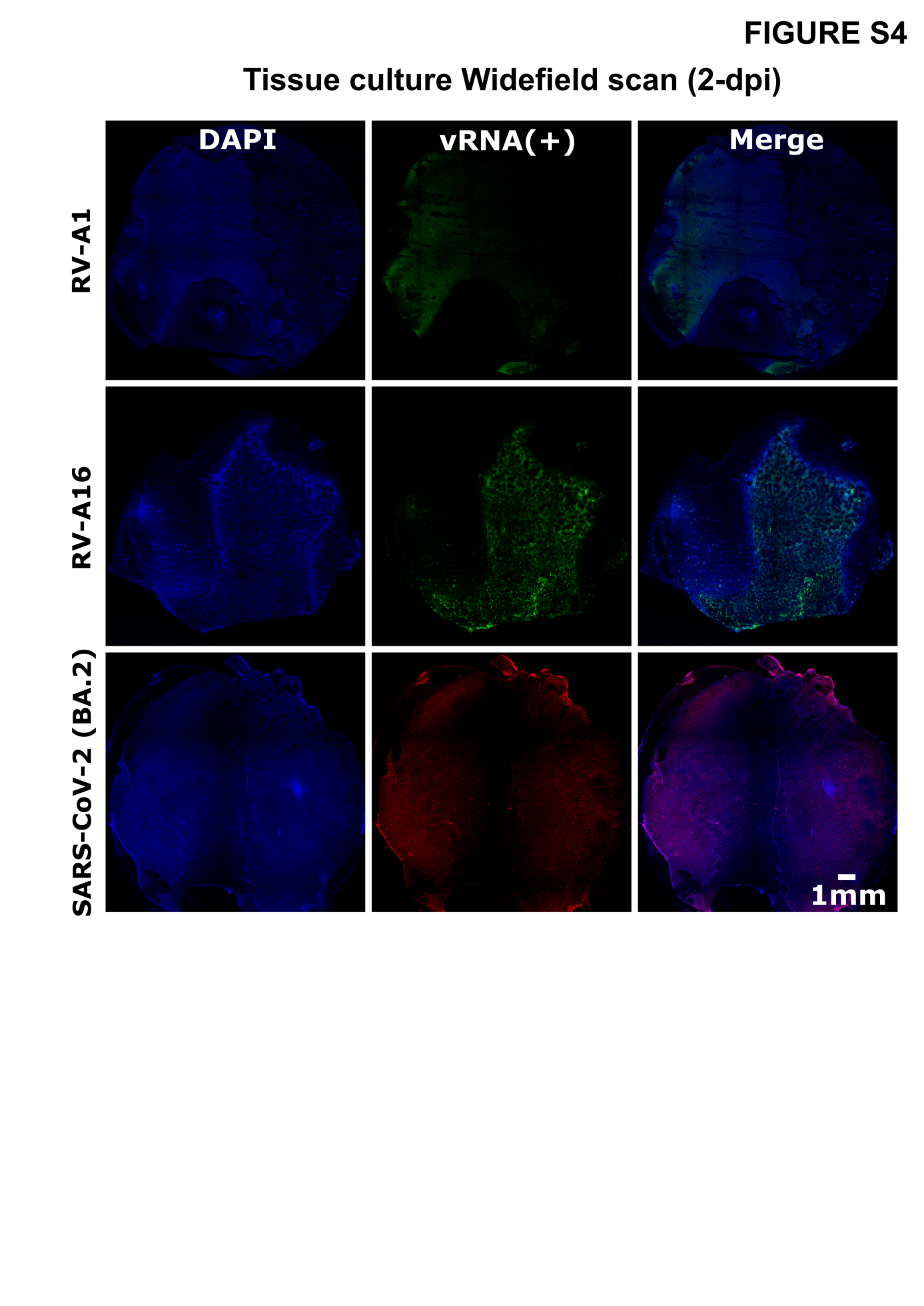

**Figure S4. Spatial distribution of SARS-CoV-2 and RVs in single tissue cultures at 2-dpi.**

Single infected bronchial epithelium were fixed at 2-days post infections and stained by RNA-FISH for SARS-CoV-2 RNA(+) ORF1ab, RV-A1 RNA(+) P3 region or A16 RNA(+) P1 region. Nuclei are stained with DAPI in blue. Tissue culture observation were done with widefield Olympus ScanR HCS microscope.
