## Supplemental Figure 5 for "DYNAMICS OF RHINOVIRUS SARS-COV-2 COINFECTIONS AND SUPERINFECTIONS IN HUMAN AIRWAY CULTURES REVEAL TYPE-DEPENDENT VIRAL INTERFERENCE"

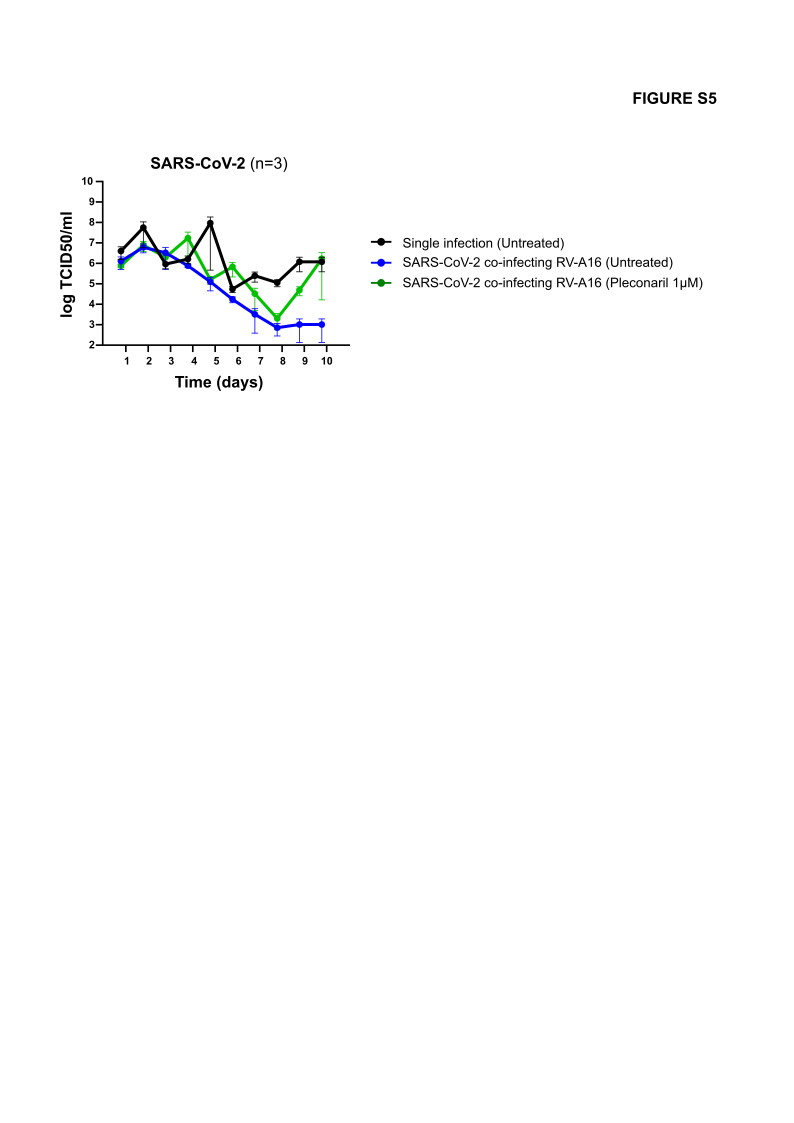

**Figure S5. Specific inhibition of RV-A16 by Pleconaril does not potentiate SARS-CoV-2 (BA.2) coinfection.** Bronchial epithelium air-liquid cultures were infected at day 0 by RV-A16 alone or with SARS-CoV-2 (10,000 TCID_50_/tissue, 33.5°C), and treated with Pleconaril (µM final) in the basolateral medium, starting at 2h post infection. Pleconaril was then administrated daily. Single RV-A16 and SARS-CoV-2 (BA.2) infections, and mock treatment were done in parallel as control. Apical SARS-CoV-2 progenies were quantified by TCID_50_ titrations on VeroE6-TMPRSS.
