## Supplemental Figure 6 for "DYNAMICS OF RHINOVIRUS SARS-COV-2 COINFECTIONS AND SUPERINFECTIONS IN HUMAN AIRWAY CULTURES REVEAL TYPE-DEPENDENT VIRAL INTERFERENCE"

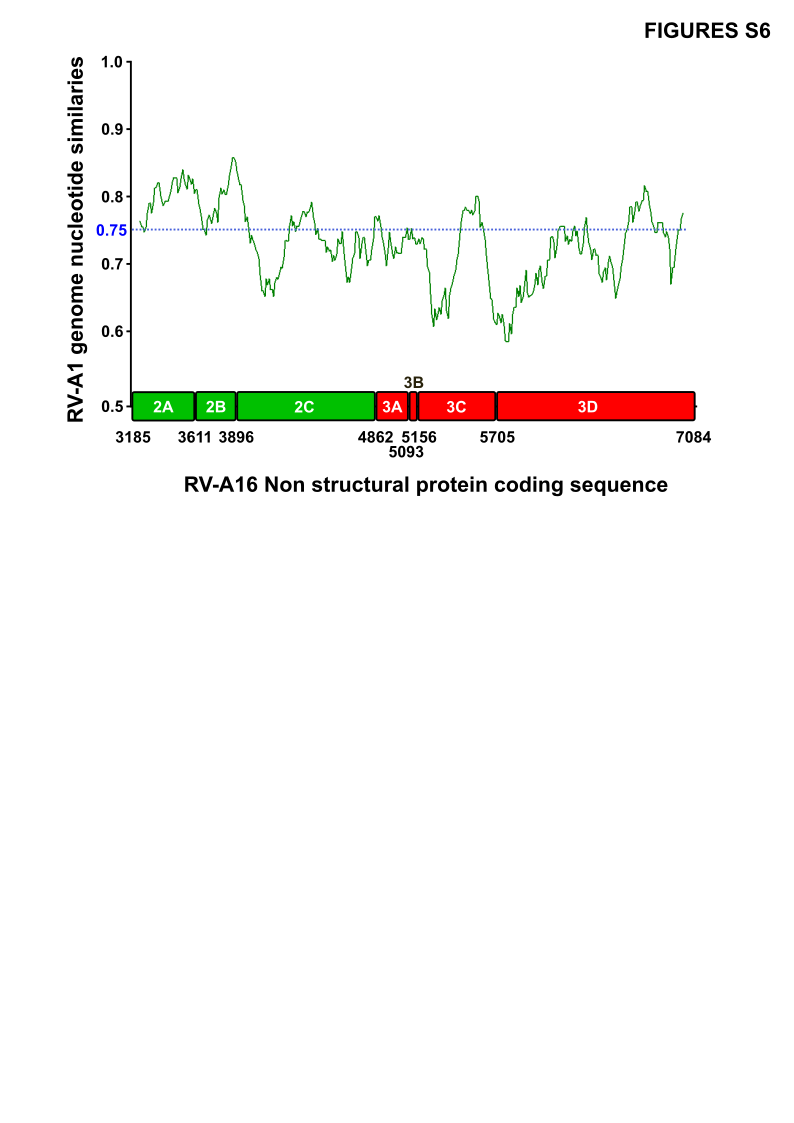

**Figure S6. Nucleotide similarity plot analyses of RV-A1 and RV-A16 non-structural coding sequences.** A scheme indicating the genomic organization of RV-A16 non-structural regions is shown and the RV-A16 sequence was used as query. Nucleotide similarity was calculated with SimPlot, version 3.5.1, software, with a 200-nt window moved along the sequence in 20-nt steps.
